# Conserved patterns of sequence diversification provide insight into the evolution of two-component systems in Enterobacteriaceae

**DOI:** 10.1101/2023.11.03.565435

**Authors:** Luke A.F. Barretto, Patryc-Khang T. Van, Casey C. Fowler

## Abstract

Two-component regulatory systems (TCS) are a major mechanism used by bacteria to sense and respond to their environments. Many of the same TCS are used by biologically diverse organisms with different regulatory needs, suggesting that the functions of TCS must adapt over evolution. To explore this topic, we analyzed the amino acid sequence divergence patterns of a large set of broadly conserved TCS across different branches of Enterobacteriaceae, a family of Gram-negative bacteria that includes biomedically important genera such as *Salmonella*, *Escherichia*, *Klebsiella*, and others. Our analysis revealed trends in how TCS sequences change across different proteins or functional domains of the TCS, and across different lineages. Based on these trends, we identified individual TCS that exhibit atypical evolutionary patterns. We observed a strong correlation for the extent of sequence variation of a given TCS across different lineages, unveiling a hierarchy TCS sequence conservation with EnvZ/OmpR as the most conserved TCS. We provide evidence that, for the most divergent of the TCS analyzed, PmrA/PmrB, different alleles were horizontally acquired by different branches of this family, and that different PmrA/PmrB sequence variants have highly divergent signal sensing domains. Collectively, this study sheds light on how TCS evolve, and serves as a compendium for how the sequences of the TCS in this family have diverged over the course of evolution.

## INTRODUCTION

All organisms face the fundamental challenge of sensing and responding to environmental changes. Two-component regulatory systems (TCS), versatile regulatory systems that couple environmental signal detection to physiological responses, are an important mechanism used by many organisms to ensure the timely expression of their genetic repertoires in response to changing conditions [1,2]. TCS can be found in plants, fungi, and archaea, but they are more prevalent in bacteria where a single genome can often encode dozens or even hundreds of different TCS [3]. The two conserved elements that define a TCS are a histidine kinase (HK), which is responsible for environmental sensing, and a response regulator (RR), which mediates a biological response. In a prototypical TCS, the HK dimerizes upon sensing stimuli and autophosphorylates by binding ATP and transferring its γ-phosphoryl group to a conserved His residue. HKs then initiate a phosphotransfer reaction to a conserved Asp residue of its cognate RR [1,4,5]. Phosphorylation of the RR leads to the rearrangement of molecular surfaces that promote dimerization of the RR and a stabilization its active conformation [4,6,7]. Many HKs are bifunctional and also act as phosphatases that inactivate their cognate RR in the absence of a stimulating signal [1,5,8]. Signal transduction through the RR can elicit a wide range of physiological responses that depend on the nature of the RR’s output domain [9]. In general, both the HK and the RR components of a TCS have a modular architecture [6,8]. A typical HK is comprised of discrete domains involved in sensing, signal transmission, and kinase activity. The kinase domain is a conserved feature of HKs and can be subdivided into a Dimerization and Histidine phosphotransfer domain (DHp) and a CAtalytic domain (CA) [8]. By contrast, the sensing and signal transduction modules are not conserved across different HKs, with sensing modules in particular exhibiting tremendous diversity both in terms of structure and the stimuli recognized [2,8,10]. Most HKs possess extracytoplasmic sensory domains (Class I), while others utilize multiple transmembrane (Class II) or even cytoplasmic (Class III) sensory domains [10,11]. RRs are composed of a conserved receiver (REC) domain, which contains the phosphorylatable Asp side chain, and a variable output domain that is responsible for eliciting biological changes in response to the phosphorylation of the REC domain by the HK. The output domain differs amongst different TCS, with DNA binding transcription factors representing the most common output modules [9,12]. Hybrid HKs elaborate on the classic TCS scheme by encoding a REC domain, and may also include Histidine Phosphotransfer (HPt) domain(s) or associated HPt proteins [1,5]. The result is a multistep phosphorelay signal transduction that shuttles the phosphoryl group from multiple His-and Asp-containing domains before phosphorylating the REC domain of the RR that elicits the biological response [5,13]. In addition to this twist on the classic TCS paradigm, there is an increasing appreciation that the regulatory cascades of many TCS involve additional components beyond the core HK/RR pair. The nature of these auxiliary components is variable, but includes a long and growing list of proteins that directly interact with the TCS (often the HK) to influence its activity level [14–16]. In many instances, these proteins serve as “connectors” that are regulated by a different regulatory system (often a different TCS) and thus allow for the integration of multiple signals into complex regulatory networks [17,18]. The diversity observed across TCS in terms of both function and architecture is the culmination of a very long and complex evolutionary history and is a testament to the efficacy of the TCS as a regulatory mechanism.

As bacteria evolve to adopt different lifestyles or to inhabit new niches, their regulatory networks must also adapt. Although TCS with a narrow phylogenetic distribution are also common, many TCS are widely distributed across evolutionarily distant taxa, indicating that the same TCS are often employed by organisms with disparate regulatory requirements. For example, the PhoP/PhoQ (PhoPQ) TCS is widely conserved amongst diverse gammaproteobacterial lineages where it senses Mg^2+^ concentrations and serves a conserved role in maintaining magnesium homeostasis [19]. However, in addition to this conserved function, PhoPQ senses a variety of other signals and controls highly variable repertoires of genes in a manner that differs amongst different lineages [19–21]. In *Salmonella enterica* (*S. enterica*), for example, PhoPQ is a master regulator that senses the unique intracellular environment that this species inhabits when it invades an animal cell, and responds by activating the expression of a battery of genes encoding proteins that range from stress response systems to toxins that disrupt host cell biology [19,21]. This intracellular niche bears little resemblance to any environment encountered by soil-dwelling or aquatic organisms that encode orthologous PhoPQ TCS. The PhoPQ regulatory network has adapted in several ways to accommodate the variable roles it must play in different organisms, including changes to its signal sensing, signal transduction and DNA binding properties, as well as differences in the repertoires and the activities of connector proteins that integrate PhoPQ signalling with other TCS [14,15,19,21–23]. The PhoPQ example is not unique, and different TCS and different organisms that encode those systems face variable evolutionary pressures depending upon their unique biology. With few exceptions, very little is known about how the functional properties of TCS differ in different organisms, and there is currently a poor overall understanding of how TCS evolve.

In this study, we explore the evolution of TCS in the Enterobacteriaceae family through an analysis of how the sequences of broadly distributed TCS have changed across this lineage. Enterobacteriaceae, a large family of Gram-negative gammaproteobacteria, is composed of a cosmopolitan assortment of organisms that inhabit diverse environments including vertebrate gastrointestinal tracts, plants, water, and soil [24]. This lineage includes predominantly environmental organisms, phytopathogens, as well as important human pathogens such as *S. enterica*, *Klebsiella pneumoniae* (*K. pneumoniae*), and assorted *Escherichia coli* (*E. coli*) pathovars, amongst others. Historically, a very broad, phylogenetically diverse assortment of bacterial lineages were assigned to Enterobacteriaceae. More recently, the Enterobacterales Order has been reorganized to yield monophyletic families that better represent the evolutionary relationships amongst the genera originally assigned to this family [25,26]. For example, phylogenetically distant species such as *Yersinia pestis* (*Y. pestis*) and *Proteus mirabilis* were historically described as Enterobacteriaceae, but now fall under the families Yersiniaceae and Morganellaceae, respectively [25,26]. We chose the Enterobacteriaceae family to investigate TCS evolution for several reasons including: (i) the importance of organisms in this taxon with regard to human health and biotechnology, (ii) TCSs are relatively well studied in this family, with species such as *E. coli* and *S. enterica* serving as model systems for the study of TCS biology, and (iii) different genera of this family are sufficiently diverse to capture tremendous phenotypic and ecological diversity that has evolved over millions of years, but sufficiently similar such that orthologous TCS can be readily identified, and that orthologous systems are likely to serve a related regulatory function in most cases. Numerous studies have explored TCS evolution through the mining and analysis of sequence databases. These studies have made many important contributions to our understanding of TCS, such as providing in-depth analyses of their phylogenetic distributions and how TCS repertoires vary across different taxa, surveying the architectural diversity of TCS and identifying conserved functional elements of HKs and RRs, providing insights into how new TCS emerge or how they are transferred between lineages, and shedding light on how HK-RR specificity is maintained over evolution [3,27–38]. In this study, we focus on the patterns of sequence diversification amongst orthologous TCS that have diverged in different genera that are separated by millions of years of evolution. Our results reveal highly variable rates of evolutionary divergence for different TCS. However, we observe strong correlations in the patterns of sequence divergence for individual TCS across different genera, or when comparing the HK and RR constituents or the functional domains that comprise these proteins. Importantly, TCS with atypical evolutionary patterns were identified, providing leads for future investigations into areas such as species-specific adaptations of individual TCS. Our analysis identified PmrA/PmrB, an important mediator of resistance to certain antimicrobial compounds, as the broadly distributed TCS with the most divergent sequence across Enterobacteriaceae, and we provide evidence that multiple different alleles of this TCS with divergent signal sensing domains were independently acquired by different branches of this family. Collectively, this study reveals how the sequences of Enterobacteriaceae’s TCS have diverged over evolutionary time and sheds new light on the general principles of how TCS evolve.

## RESULTS

### Analysis of the evolutionary relationships between the selected Enterobacteriaceae reference strains and their TCS repertoires

To analyze the evolution of TCS in Enterobacteriaceae, we first generated a set of reference genomes. We selected one strain from each of nine different genera within this family: *Escherichia coli* K-12 (strain MG1655), *Salmonella enterica* (serovar Typhimurium, strain LT2), *Citrobacter rodentium* (strain ICC168), *Klebsiella variicola* (strain 342; originally assigned to the *K. pneumoniae* species but later reassigned to the *K. variicola* species, which is a member of the *K. pneumoniae* complex), *Cronobacter turicensis* (strain z3032), *Enterobacter cloacae* (strain ATCC 13047), *Phytobacter diazotrophicus* (strain TA9730), *Kosakonia sacchari* (strain BO-1), and *Huaxiibacter chinensis* (strain ZB04; originally referred to as *Lelliottia amnigena*, but later assigned to *H. chinensis*) [39,40]. The goal of this reference strain set was not to comprehensively capture the genetic diversity within this lineage, but rather to serve as a cross-section of Enterobacteriaceae TCS genetic diversity, such that individual strains represent genera that have diverged in ecology and lifestyle over millions of years. To provide a framework for exploring the evolutionary relationships of the TCS encoded by these strains, we generated a phylogenetic tree using *Y. pestis* strain CO92 as an outgroup. This tree was generated based on multiple sequence alignments (MSA) of the amino acid (AA) sequences of the proteins encoded by the core genome common to all reference strains (Fig. 1A). The results of this analysis, which are congruent with previous analyses of Enterobacteriaceae phylogenetics, confirm that the reference strains selected are well dispersed across this family [25,41]. Although well-dispersed, certain lineages are predicted to have diverged from one another more recently, such as *E. cloacae*/*H. chinensis* or the *E. coli*/*S. enterica*/*C. rodentium* clade. By contrast, *C. turicensis* is predicted to have branched off from the other lineages relatively early in the evolutionary history of this family.

**Figure 1:**
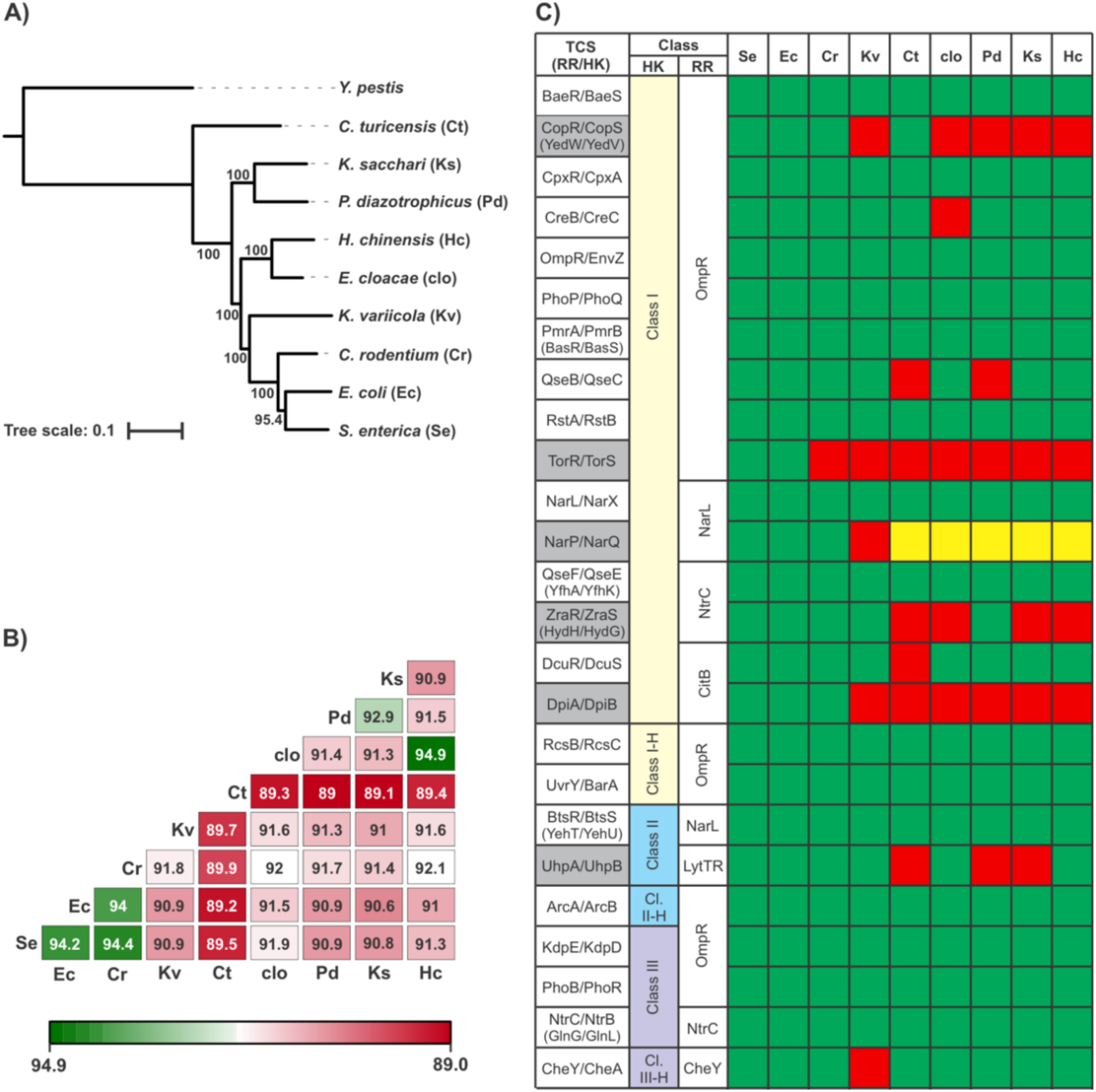
Overview and analysis of the reference strains and TCS analyzed in this study. **(A)** Phylogenetic tree of the reference strains selected for analysis in this study. The evolutionary relationships between the 9 chosen reference strains, each from a different major genus of the Enterobacteriaceae family, were analyzed by generating a phylogenetic tree based on the amino acid sequence alignments of the proteins encoded by the core genome of these strains. *Y. pestis* strain CO92, a member of the Enterobacterales order but not the Enterobacteriaceae family, was included as an outgroup. Numbers represent bootstrap support values calculated from 500 replicates. Lineages are listed as species; strain names are provided in the main text. **(B)** Matrix of the average AA sequence identities of the conserved gene set between all pairwise combinations of reference strains. Numbers indicate the average percent AA sequence identity across the 50 selected conserved genes; matrix is shown as a heat map for visual clarity. **(C)** The 25 TCSs conserved in *E. coli* and *S. enterica* were analyzed for their distribution amongst the reference strains set, where green indicates the presence of both the RR and HK, yellow indicates the absence of one component, and red denotes the absence of both the HK and the RR. TCSs that were absent in more than 2 reference strains (and not selected for further analysis) are shaded in grey. TCS are classified based on their architecture and the nature of their output domain. HK Architectures (HK) are designated as either: Class I (yellow), the classic HK archetype, which feature a periplasmic-sensing domain linked by a transmembrane domain to its cytoplasmic kinase domain, Class II (Blue) which use multiple transmembrane regions for sensing but have a cytoplasmic kinase domain, or Class III (Purple) which sense cytoplasmic signals and can be either membrane-associated or soluble [10]. TCS were further classified according to the output domain of their RR. Other than CheY, which lacks a discrete output domain, all other RRs possess a DNA-binding output domain belonging to one of five families: OmpR, NarL, NtrC, CitB, and LytTR [9,12]. Shorthand used for strains in all panels (e.g. “Ec”) is described in panel (A).

For certain analyses presented below, we sought to contextualize the evolution of TCS proteins by comparing their divergence to that of other proteins that are conserved within Enterobacteriaceae. To this end, we assembled a list of 50 genes that were present in all reference strains and that have highly conserved functions spread across assorted functional categories (Table S1). We compiled the amino acid (AA) sequences of these 50 conserved genes (CGs), conducted multiple sequence alignments (MSAs) for each protein, and generated percent AA sequence identity matrices for each CG across the reference strains, as well as a cumulative matrix that represents the average values for the 50 proteins (Fig 1B, Supplemental Datasets 1-2). The phylogenetic distances between reference strains inferred by the cumulative matrix are consistent with the phylogenetic tree generated above, and with previously published results (Fig. 1A-B) [25,42]. The average AA sequence identity of the 50 CGs between species ranges from ∼89% (*C. turicensis* compared all other species) to ∼94% (*E. coli*/*S. enterica*/*C. rodentium*, as well as *E. cloacae*/*H. chinensis*); these values are somewhat higher than genome-wide averages reported previously comparing some similar combinations of strains, which was expected given that our gene set was limited to highly conserved genes with integral functions [41,43,44]. The narrow spread in average AA sequence identities across all combinations of reference strains indicates that this collection of strains represents a well-dispersed cross section of the Enterobacteriaceae family (Fig 1B). When the results of our phylogenetic and CG sequence analyses are contextualized using previously reported estimates of the dates that some combinations of these species diverged from a common ancestor, it is likely that all of our reference strains are separated by ∼100 million of years of evolution or more [45,46]

We next compiled the set of TCS that are broadly conserved amongst our reference strains. Since *E. coli* and *S. enterica* have served as important model organisms for studying TCS biology, we first generated a complete list of HKs and RRs from the genomes of these two reference strains. Because we sought to conduct global analyses of how HK and RR pairs have diverged over evolution, we limited our list to cognate HK/RR pairs, and disregarded TCS proteins that did not fit this mold, such as orphan HKs/RRs without overt interacting partners, or auxiliary proteins which expand the TCS phosphorelay. We found that 25 TCS are conserved in both species, which is in good agreement with previous studies (Fig. 1C) [3,47,48].

Extracytoplasmic Class I HKs dominated the set of conserved TCS, followed by cytoplasmic Class III HKs and transmembrane Class II HKs (Fig 1C). With the exception of the chemotaxis RR CheY (which lacks an output domain), all other RRs have DNA-binding output domains, which is notable given that DNA-binding output domains comprise only ∼66% of functional RR subfamilies identified in Pfam [12]. The nature of the DNA binding domains in the 25 RRs largely mirrors the overall prevalence of DNA-binding RR subfamilies in sequence databases (Pfam), with OmpR being the most common, followed by NarL, NtrC, CitB, and LytTR [12]. We then analyzed the TCS list generated based on *E. coli*/*S. enterica* for their distribution across the other seven reference strains (Fig 1C). We found that, in general, TCS were broadly conserved across the family, and that the majority of TCS were found in most or all of the reference strains. Although we aimed to select a broad assortment of TCSs for further analysis, the inclusion of TCS with a narrow phylogenetic distribution would potentially skew downstream analyses that examine TCS evolutionary trends. To balance these considerations, we omitted any TCS that was absent from more than two of the nine reference strains (TCS names shaded grey in Fig 1C); this cutoff eliminated 6 of the originally identified 25 TCS, and the remaining 19 broadly conserved TCS were selected for further analysis.

### Population-level analysis of TCS sequence diversification in Enterobacteriaceae

To begin our investigations into the evolutionary diversification of TCS in Enterobacteriaceae, we compiled the amino acid sequences of the 19 selected TCS in each of our reference strains and conducted MSAs for each HK and RR (Table S2, Supplemental Datasets 3-4). To facilitate the analysis of these data, we generated a matrix of the AA divergence rates for each HK and RR (which we define as the number of AA sequence differences per 100 AAs) between each possible pairwise combination of the nine reference strains (Supplemental Dataset 2). Using these data, we then determined the average rates at which RRs and HKs accumulated amino differences across our entire dataset, and how these values compared to the average rates observed for our CG dataset (Fig 2A). We found that, when averaged across all proteins analyzed and all pairwise species comparisons, RRs exhibited 9.9 AA differences/100 AAs, a very similar value to the 8.7 AA differences/100 AA observed for our CG set. This suggests that, at a population level, the AA sequences of broadly conserved RRs have diverged at a similar rate as a typical broadly conserved protein within Enterobacteriaceae. By contrast, we found that HKs had a significantly greater divergence rate (p <0.01 when compared to CGs) of 17.9 AA differences/100 AAs, indicating that, on average, HKs accumulate sequence changes at about double the rate of RRs and of conserved proteins at large. Given that RRs and HKs generally work together as a single functional unit, the substantially greater rate of sequence change in HKs compared to RRs suggests that evolutionary adaptations to the functional properties of a TCS might be more common in the sensing and signal transduction component than in the output protein.

**Figure 2:**
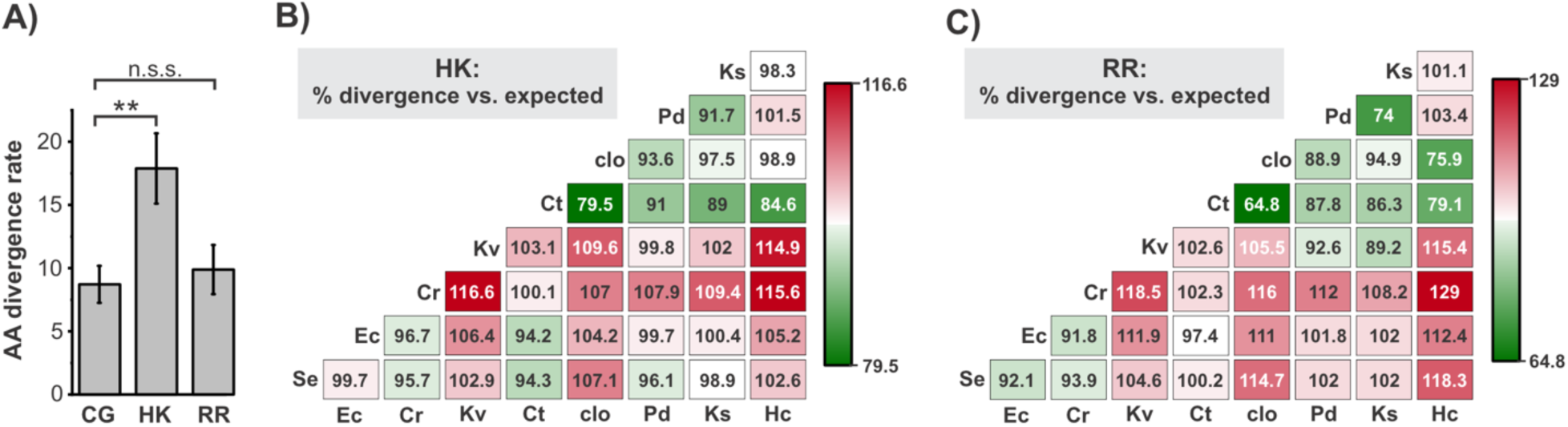
Population-level trends in TCS divergence across Enterobacteriaceae. **(A)** The average divergence rate (AA differences/100 amino acids) across all proteins analyzed and all pairwise combinations of reference strains for HKs, RRs, and the conserved gene (CG) set. Error bars represent the standard deviation. The statistical significance of the differences in divergence rates of HK and RR compared to the CG was analyzed using a one-way ANOVA test; ** indicates p < 0.001, n.s.s. indicates not statistically significant. **(B and C)** Matrix showing the divergence rates averaged across all HKs (B) or RRs (C) compared to the expected value for all combinations of the reference strains. The expected average divergence rate for a given pair of strains was calculated by multiplying the average divergence rate for those strains across the CG set by a correction factor that accounts for differences in the average divergence rate of CG compared to HKs (C) or RRs (D). The divergence rate compared to expected is expressed as a percentage, such that values exceeding 100% represent pairs of reference strains whose TCS proteins have diverged more than expected. Matrix is presented as a heat map for visual clarity.

We next examined the patterns of sequence change for TCS across the different combinations of reference strains and how this compared to the patterns observed for our CG set. Because TCS are a central mechanism by which Enterobacteriaceae sense and respond to their environment, we reasoned that, at the population level, the TCS of species that have adapted to very different niches might be disproportionately divergent (i.e. more divergent on average than the CGs, which have diverse functions that do not necessarily relate to niche adaptation). To explore this possibility, we generated matrices of the expected divergence rates for both RRs and HKs across the reference strains, which were derived using the average AA divergence rates of the CGs, multiplied by a correction factor that accounts for the observed differences in average divergence rates of CGs/HKs/RRs (Fig 2B). We then compared these expected values to the observed average divergence values for HKs and RRs to generate matrices of the AA divergence compared to expected (Fig 2B-C). We found that the HK and RR matrices showed strikingly similar trends across the various species comparisons, indicating a strong correlation in the extent of sequence divergence of the two constituents of TCS across Enterobacteriaceae. Although various combinations of organisms exhibited more or less TCS sequence variation than expected, certain trends were apparent in these data. For example, more variation than expected was observed for 13 of the 16 comparisons involving *K. variicola* HKs/RRs, suggesting that TCS on the whole are unusually divergent in this lineage. This is consistent with the remarkable ecological diversity of *Klebsiella* species, which inhabit a very wide range of environments, including assorted free-living and host-associated niches [49,50]. We also found that the TCS of the *E. coli*/*S. enterica*/*C. rodentium* clade are somewhat more similar to one another than expected, but they are almost universally more divergent than expected when compared to other lineages. This is in agreement with the adaptation of this clade to animal intestinal tracts, which is not thought to be the predominant environmental reservoir for the other lineages. Interestingly, many of the pairwise comparisons yielding the lowest rates of TCS divergence compared to expected involved the *Cronobacter*/*Enterobacter*/*Phytobacter/Kosakonia* lineages (Fig 2B-C). Although little is known about the natural ecology of these lineages and they can be isolated from various environmental niches such as soil or water, it is noteworthy that numerous strains amongst these genera have been identified as plant-associated or as plant pathogens [24,51–57]. The overall trend, therefore, was that TCS were more divergent than expected when comparing species adapted to animal hosts to those adapted to environmental niches/plant hosts, but less divergent than expected when comparing within those groups; the TCS of the ecologically flexible *Klebsiella* lineage was divergent from both groups. Collectively, these data indicate that the extent to which the sequences of TCS diverge over evolution is disproportionately dictated by the nature of the niches they inhabit when compared to the proteome at large.

### Sequence divergence patterns of individual TCS across Enterobacteriaceae

We next examined the evolutionary patterns of individual TCS. First, we compared the mean rates at which individual TCS proteins have diverged across Enterobacteriaceae by averaging the divergence rates from all of the possible pairwise combinations of our reference strains (Table 1). Consistent with previous observations that have noted coevolution for TCS constituents, we observed a strong positive correlation in the divergence rates of HK and RR partners (Fig 3A) [27]. However, despite the clear co-evolutionary trend across all TCS, there is variability in this relationship from TCS to TCS. Above, we noted that the divergence rates of HK are, on average, ∼2-fold higher for HK than for RR. Although this ratio is in this range for many HK-RR pairs, several TCS exhibit ratios as low as ∼1 (where the HK and RR have diverged at a similar rate), while for other HK-RR pairs this ratio is as high as ∼4 (Fig 3B). As an example to illustrate this variability, the HK component of the RcsBC TCS has an average divergence rate of 16.3 AA differences/100 AA, which is ∼40% higher than that of the BtsRS TCS (11.6 AA differences/100 AA). By contrast, the BtsRS RR is more variable than an average RR (12 AA differences/100 AA), while the RcsBC RR is amongst the most highly conserved of all RR (2.9 AA differences/100 AA). The factors underlying the relative variability of the HK and RR components of a given TCS are presumably multifaceted. In the case of the RcsBC TCS, the complex architecture of this signalling cascade might be a relevant factor in its atypical HK:RR divergence ratio [58]. Overall, these data indicate that HK-RR pairs co-evolve, but that the relative rates of HK and RR divergence varies amongst different TCS.

**Figure 3:**
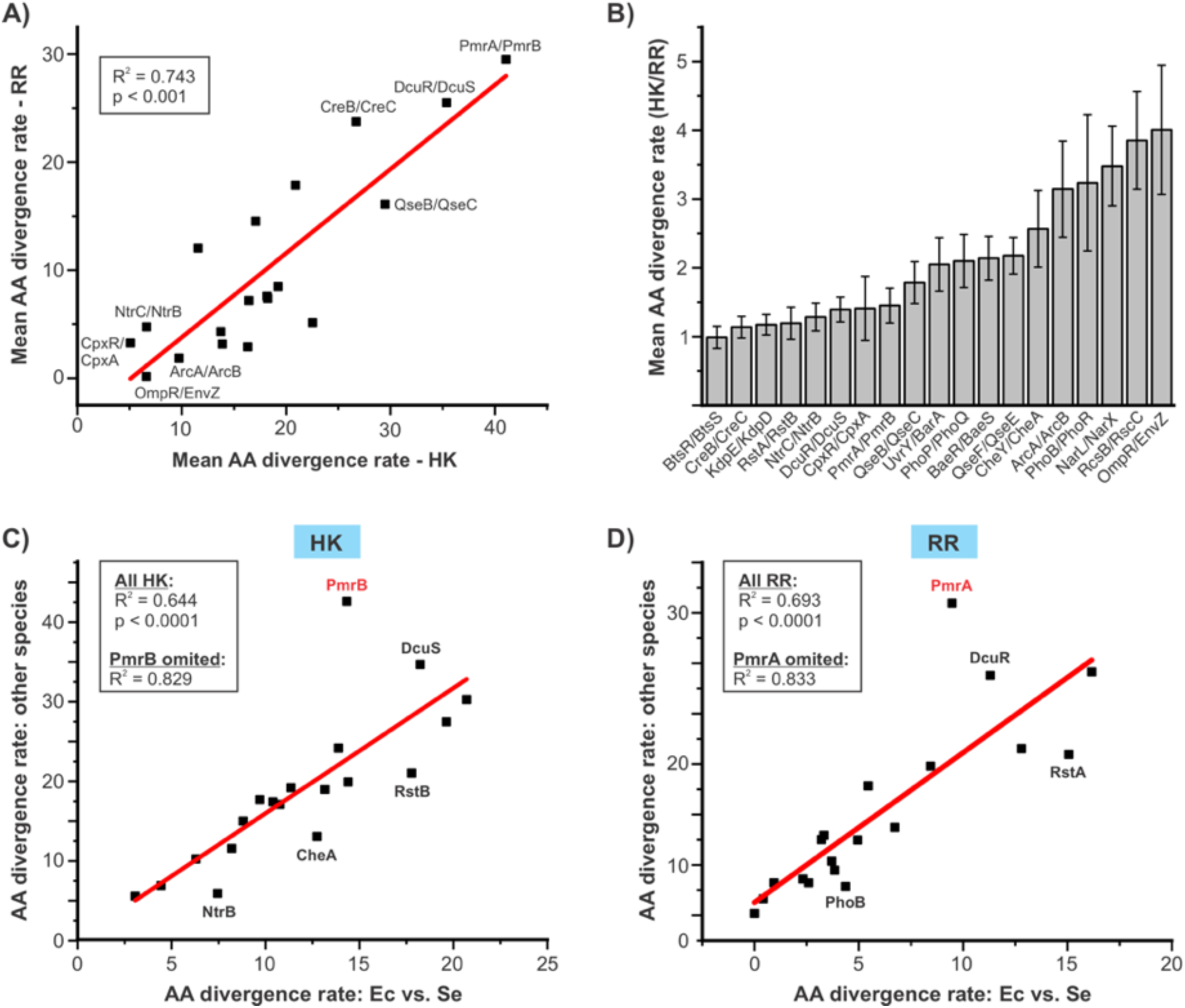
Trends in the sequence divergence of individual TCS across Enterobacteriaceae. **(A)** There is a strong positive correlation in the AA sequence divergence rates of the HK and RR constituents of a TCS. For each TCS, the average divergence rates across all pairwise combinations of reference strains were calculated for its HK and RR. Scatterplot shows the average divergence rate (AA differences/100 AA) for the HK compared to its cognate RR. The identities of select TCS with atypically low or high divergence rates are shown. **(B)** Bar graph showing the ratio of the average divergences rates of each HK compared to its cognate RR. **(C and D)** The extent to which TCS diverge in one branch of Enterobacteriaceae strongly correlates with its divergence in independent branches of this family. For each HK (C) and RR (D) the divergence rate when comparing its *E. coli* AA sequence to its *S. enterica* sequence was compared to its average divergence rate across all pairwise combinations of the other seven reference strains using scatterplots. Select individual HKs or RRs that deviate from the strong correlation observed between these two variables are identified. PmrA and PmrB (red text) were both determined to be statistical outliers in these analyses, which is shown in Fig S1. A similar analysis using *E. cloacae*/*H. chinensis* as the reference comparison is also shown in Fig S1. For all scatterplots, p values (shown within plots) were derived from the linear regression analysis and indicate highly significant positive correlations (i.e. that the correlation coefficients are greater than zero).

**Table 1.** List of Two component systems (TCSs) analyzed in this study, from least divergent to most.

| TCS (RR/HK) | Number of Species<sup>a</sup> | Species Absent | HK Divergence rate<sup>b</sup> | RR Divergence rate<sup>b</sup> | Average (HK & RR) |
| --- | --- | --- | --- | --- | --- |
| OmpR/EnvZ | 9 |  | 6.6 | 0.2 | 3.4 |
| CpxR/CpxA | 9 |  | 5.1 | 3.3 | 4.2 |
| NtrC/NtrB | 9 |  | 6.6 | 4.7 | 5.7 |
| ArcA/ArcB | 9 |  | 9.7 | 1.9 | 5.8 |
| PhoB/PhoR | 9 |  | 13.9 | 3.2 | 8.5 |
| CheY/CheA | 8 | *K. variicola* | 13.7 | 4.3 | 9.0 |
| RcsB/RcsC | 9 |  | 16.3 | 2.9 | 9.6 |
| BtsR/BtsS | 9 |  | 11.6 | 12.0 | 11.8 |
| UvrY/BarA | 9 |  | 16.4 | 7.2 | 11.8 |
| QseF/QseE | 9 |  | 18.2 | 7.4 | 12.8 |
| BaeR/BaeS | 9 |  | 18.2 | 7.6 | 12.9 |
| NarL/NarX | 9 |  | 22.5 | 5.1 | 13.8 |
| PhoP/PhoQ | 9 |  | 19.2 | 8.5 | 13.9 |
| KdpE/KdpD | 9 |  | 17.1 | 14.5 | 15.8 |
| RstA/RstB | 9 |  | 20.9 | 17.9 | 19.4 |
| QseB/QseC | 7 | *C. turicensis*<br>*P. diazotrophicus* | 29.5 | 16.1 | 22.8 |
| CreB/CreC | 8 | *E. cloacae* | 26.7 | 23.8 | 25.2 |
| DcuR/DcuS | 8 | *C. turicensis* | 35.4 | 25.5 | 30.4 |
| PmrA/PmrB | 9 |  | 41.1 | 29.5 | 35.3 |

<sup>a</sup> Denotes the number of species in the reference strain set (out of a total of 9) in which the TCS is present
<sup>b</sup> Divergence rate indicates the average number of AA differences per 100 AAs in all pairwise comparisons of HKs/RRs among all the species in which the HK/RR is present

The most overt observation from our analysis of individual TCS was that the average divergence rates varied drastically from TCS to TCS (Table 1). The most conserved TCS protein, the RR OmpR, was found to be more highly conserved than any of the 50 proteins in the CG set, with a remarkably low divergence rate of <0.2 AA differences/100 AA. By contrast, the most divergent protein was the HK PmrB with a divergence rate of 41.1 AA differences/100 AA; this rate is more than 200-fold higher than that of OmpR, nearly five-fold higher than the CG average, and greater than any of the proteins in the CG set. Notably, the extent to which a given TCS’s sequences were conserved or variable tended be consistent across different species comparisons (Supplemental Dataset 2). To explore this phenomenon, we plotted the divergence rate for each HK (Fig 3C) and RR (Fig 3D) for the *E. coli / S. enterica* comparison against its average divergence rate amongst all reference strain combinations that do not involve these two species. Because *S. enterica* and *E. coli* diverged after the other seven reference strains branched off from one another (Fig 1A), any sequence differences between these two strains should be independent of any differences between the other reference strains. Despite this, for both HKs and RRs, we observed a strong positive correlation (R^2^ = 0.644/0.694 for HK/RR, p < 0.0001 for both) between their divergence rates in *E. coli* and *S. enterica* and the average divergence rates amongst the other reference strains. The spread of these data indicates that a surprisingly high proportion of TCS proteins fit very well to the linear regression, but that a few TCS deviate from this relationship. This indicates that, in most cases, the extent to which a given TCS has diverged between *E. coli* and *S. enterica* is predictive of how much it has diverged across independent branches of the Enterobacteriaceae family. A notable TCS for which this relationship differs is PmrAB, which was a statistical outlier for the both the HK and the RR analyses (Fig 3 C-D, Fig S1). In fact, removing PmrAB from this analysis increases the correlation coefficient markedly (R^2^ > 0.8) for both HKs and RRs (Fig 3C-D, S1). PmrAB’s unusual pattern of evolution is explored further in subsequent analyses presented below. The predictive nature of TCS divergence between different Enterobacteriaceae lineages is not an anomaly restricted to *E. coli*/*S. enterica* since the same analysis using *E. cloacae*/*H. chinensis* as the reference comparison yielded a similarly strong correlation (Fig S1). This suggests that the dominant factor driving the evolutionary divergence rates of individual TCS in our dataset is the nature/function of the TCS itself, rather than the unique biological features or evolutionary pressures of the various organisms.

### Species-specific evolutionary adaptations of conserved TCS

The analyses above suggest that TCS divergence rates are correlated amongst different lineages, but also that certain TCS proteins deviate from this relationship. For example, the extent to which the PmrAB, DcuRS and RstAB TCS have diverged between *E. coli* and *S. enterica* differs from what would be expected based on their divergence rates in other Enterobacteriaceae (Fig 3C-D). To identify TCS whose sequences have changed disproportionately in a single reference strain, we analyzed how much each TCS protein has diverged in each reference strain compared to what would be expected based on its average divergence rate across the other reference strains (Table 2, Supplemental Dataset 5). These results show that, for the vast majority of TCS proteins in all reference strains, their sequence has diverged at a rate that is similar to the expected value, which is consistent with the strong predictive trend noted above. However, certain TCS proteins have diverged significantly more than expected in select organisms, which might be indicative of TCS functional adaptations unique to that species (Table 2). One interesting example of this is the RstAB TCS, which was generally more variable than expected throughout the *E. coli*/*S. enterica*/*C. rodentium* clade (Supplemental Dataset 5). Although the function of the RstAB is not well understood, in these lineages it has been observed that this TCS plays a variety of regulatory roles related to virulence, and that its expression is regulated by the prominent virulence regulator PhoPQ [59–61]. This suggests that the RstAB TCS might have taken on novel roles that relate to animal pathogenesis in these lineages, triggering the need for functional adaptions. In *S. enterica,* RstA, the RR, is amongst the most divergent proteins compared to expected, but its cognate HK, RstB, is not (Table 2, Supplemental Dataset 5). This is noteworthy because there appear to be significant differences in RstA’s regulon in *S. enterica* and *E. coli*, and, intriguingly, RstA has been shown to induce the expression of iron uptake machinery in *S. enterica* in a PhoPQ-dependent, but PmrB-independent manner [62–64]. Coupled with the data presented here, this suggests that PmrA might have adapted to fill a novel virulence role that is independent of its cognate HK in *S. enterica*.

**Table 2.** List of response regulators (RRs) and histidine kinases (HKs) whose sequences have diverged significantly more in a particular reference strain than expected.

| Species | RR |  |  | HK |  |  |
| --- | --- | --- | --- | --- | --- | --- |
|  | Protein | Percent of Expected<sup>a</sup> | Significance<sup>b</sup> | Protein | Percent of Expected<sup>a</sup> | Significance<sup>b</sup> |
| *S. enterica* | RstA<br>NtrC | 128<br>118 | $p < 0.01$<br>$p < 0.01$ | NtrB | 135 | $p < 10^{-4}$ |
| *E. coli* | PhoB | 143 | $p < 10^{-4}$ | CheA<br>BtsS<br>RstB | 123<br>113<br>110 | $p < 10^{-4}$<br>$p < 0.01$<br>$p < 0.05$ |
| *C. rodentium* | RstA<br>QseB | 137<br>113 | $p < 10^{-4}$<br>$p < 10^{-3}$ | KdpD<br>RstB<br>CheA | 123<br>115<br>113 | $p < 10^{-7}$<br>$p < 10^{-3}$<br>$p < 0.05$ |
| *K. variicola* | CpxR<br>QseF<br>NtrC<br>BtsR<br>KdpE | 127<br>123<br>113<br>110<br>110 | $p < 0.05$<br>$p < 10^{-4}$<br>$p < 0.05$<br>$p < 0.05$<br>$p < 0.05$ | RcsC<br>QseE<br>EnvZ<br>BarA<br>ArcB | 122<br>122<br>115<br>114<br>110 | $p < 10^{-9}$<br>$p < 10^{-8}$<br>$p < 0.01$<br>$p < 10^{-4}$<br>$p < 10^{-3}$ |
| *C. turicensis* | OmpR<br>PhoP<br>BaeR<br>BtsR<br>NarL | 185<br>126<br>119<br>118<br>117 | $P < 0.05$<br>$p < 10^{-5}$<br>$p < 10^{-3}$<br>$p < 10^{-4}$<br>$p < 0.05$ | BtsS<br>NarX<br>CpxA | 112<br>110<br>110 | $p < 0.01$<br>$p < 10^{-3}$<br>$p < 0.05$ |
| *E. cloacae* | OmpR<br>PmrA | 255<br>124 | $p < 10^{-4}$<br>$p < 0.05$ | | | |
| *P. diazotrophicus* | UvrY | 115 | $p < 0.01$ | | | |
| *K. sacchari* | | | | BaeS<br>EnvZ | 113<br>111 | $p < 10^{-3}$<br>$p < 0.05$ |
| *H. chinensis* | | | | CpxA | 114 | $p < 0.01$ |

<sup>a</sup> Percent of expected indicates how much the sequence of that TCS protein in that reference strain deviates from its sequence in the other reference strains compared to what would be expected based on sequence divergence patterns in the CG set and how variable the sequence of that TCS protein is on average. Only TCS proteins with a value of at least 110% of expected are listed. See methods for more information.
<sup>b</sup> Only TCS proteins with a $p < 0.05$ are listed. See methods for more information.

### Functional domain-level analysis of TCS evolution

We next explored the evolutionary diversification of Enterobacteriaceae TCS at the level of their functional domains. This analysis was conducted to determine to what extent the strong correlation in the sequence diversification of the RR and HK constituents of a given TCS would hold true when comparing the individual functional domains. We also reasoned that these analyses could shed light on the evolutionary trajectories of individual TCS. In all TCS, the HK contains a highly conserved kinase domain, and the RR contains a highly conserved receiver domain, but other segments of these proteins are variable. To analyze the functionally and architecturally diverse TCS in our dataset in a consistent manner, we partitioned their proteins as follows. For RRs, the receiver (REC) domain boundaries were identified (see methods), and the downstream DNA binding domain (DBD) was defined as the complete sequence that was C-terminal of this sequence. The DBD, as defined here, therefore also includes the short linker that typically connects the REC and DBD of a RR. HKs were also broken into two domains: (i) the conserved kinase domain spanning from the start of the DHp domain through the end of the CA domain, and (ii) the N-terminal segment of the HK, consisting of everything prior to the first AA of the kinase domain. The architecture and function of this N-terminal segment is variable in different TCS. In most HKs, however, this region consists of sensing, signal transduction and transmembrane motifs. For simplicity, we refer to this region as the sensing domain in accordance with its predominant function. Because of substantial differences in the architecture of the chemotaxis TCS CheY/CheA relative to the others, it was omitted from this analysis [65]. We performed MSAs for each domain of each TCS protein across our reference strains and generated AA divergence rate matrices (Supplemental Dataset 6); for each TCS, the average divergence rates for each of its constituent domains are shown in Fig. 4A. We further analyzed the relationship between the divergence rates of each of the six possible combinations for the four functional domains (Fig 4B-G).

**Figure 4:**
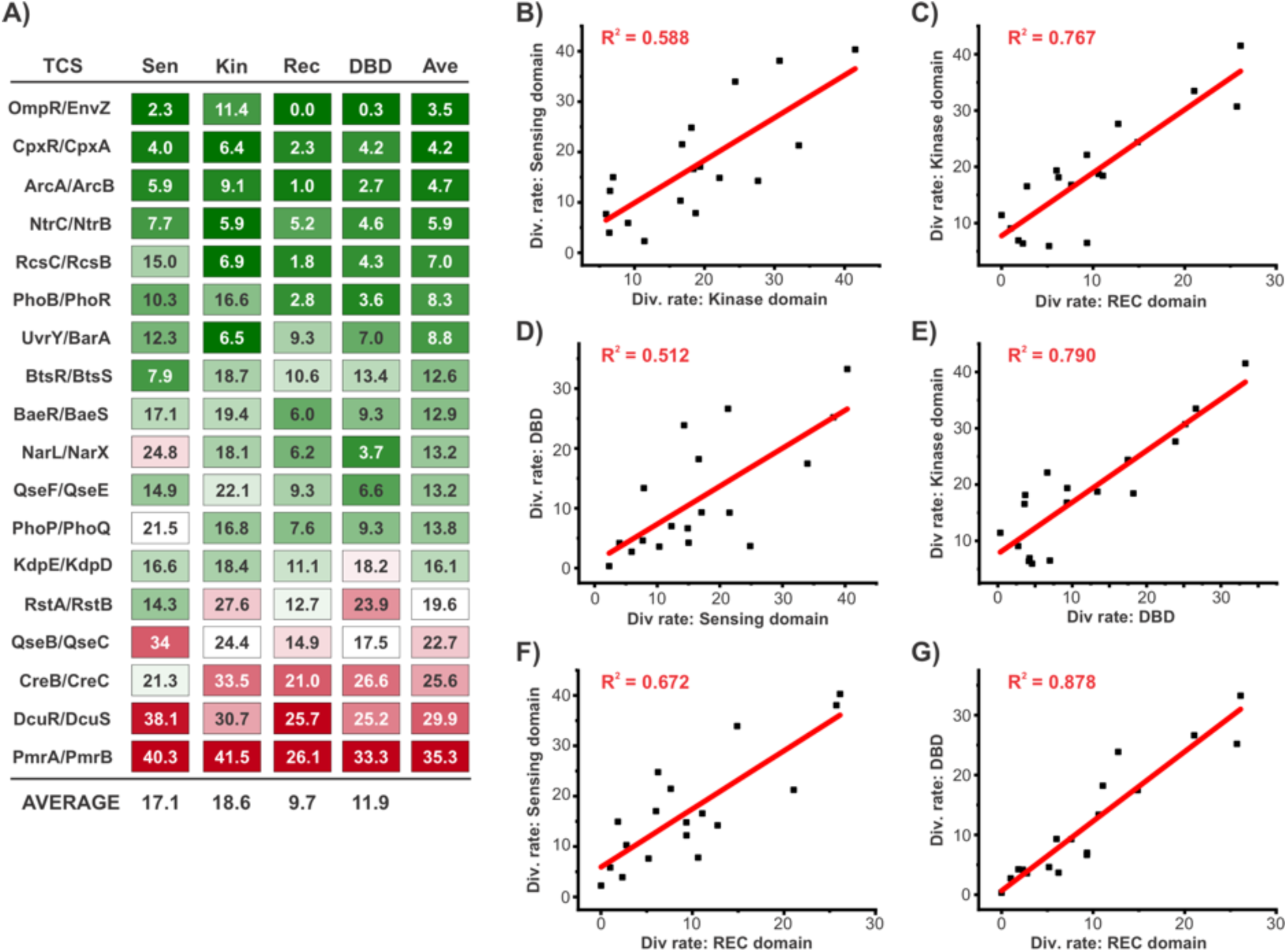
Domain-level analysis of TCS sequence divergence in Enterobacteriaceae. **(A)** Average divergence rates of individual functional domains for each TCS across the reference strains. For both the HK and the RR component of each TCS, two major functional domains were identified as outlined in the main text; sensor (Sen) and kinase (Kin) domains for the HK, and receiver (REC) and DNA binding domains (DBD) for the RR. Heat map chart shows the average divergence rate (AA differences/100 AA) across all pairwise combinations of reference strains. “Ave” indicates the average value for the four domains of that TCS. **(B – G)** Scatterplots of the average divergence rate for each TCS comparing one functional domain [as described in (A)] to another in a pairwise fashion. Each plot represents a different combination of domains, as indicated. For all scatterplots, linear regression analysis yielded a p < 0.001 that the correlation coefficient was greater than zero, indicating a statistically significant positive correlation.

Overall, the kinase domain was observed to be the most variable on average (18.6 AA differences/100 AA), followed by the sensor domain (17.1 AA differences/100 AA) and DBD (11.9 AA differences/100 AA), while the REC domain was the most highly conserved (9.7 AA differences/100 AA). The divergence rates of all pairwise combinations of domains exhibited a statistically significant positive correlation, but the strength of this correlation varied substantially for different combinations, with R^2^ values ranging from slightly above 0.5 to nearly 0.9 (Fig 4B-G). Despite the kinase domain being the most variable overall, the sensor domain exhibited the weakest correlations with the others, which is likely driven (at least in part) by different sensor domains diverging at different rates due to their architectural variability. Given that the sensor and kinase domain are both part of the HK and are functionally interconnected, we found it interesting that the divergence rate of the sensor domain correlated more strongly with the REC domain of its cognate RR, than it did to its associated kinase domain. Despite the population-level correlations for all domains, inspection of the divergence rates for individual TCS reveals substantial variability from TCS to TCS (Fig 4A). For example, for RstAB, the divergence rate of its DBD is nearly twice as high its sensor domain, which is highly atypical amongst the complete set of TCS and suggests that the output functions of this TCS might have changed more than its sensory functions. By contrast, for NarLX, the reverse situation was observed, where the divergence rate of its sensor domain is nearly 7-fold higher than that of its DBD. This suggests that the output functions of its RR are likely to be highly conserved, but the signal sensing functions of the HK might have adapted in different species. Another interesting example is the EnvZ/OmpR TCS, where the REC, DBD and sensor domains are all easily the most highly conserved of any TCS with divergence rates between 0 and 2.3 AA differences/100 AA. Given this remarkable sequence conservation, it is noteworthy that its kinase domain has a markedly higher divergence rate of 11.4 AA differences/100 AA, suggesting that any species-specific functional adaptations in this important TCS are likely to map to the cytoplasmic portion of its HK. Although the predominant function of this domain is generally signal transduction and the phosphorylation/dephosphorylation of the RR, for EnvZ it has been shown that the cytoplasmic region (“kinase domain”) is also directly involved in signal sensing, suggesting that the variability we observe here could be reflected in altered signal transduction, signal sensing, or both [66].

### Multiple sequence variants of the PmrAB TCS appear to have been independently acquired by different branches of the Enterobacteriaceae Family

PmrAB is a canonical TCS composed of the class I HK PmrB and the RR PmrA, which is an OmpR-family transcription factor [67]. PmrB’s periplasmic sensory domain detects signals that include high concentrations of Fe^3+^ and a mildly acidic pH, which promote PmrB’s phosphorylation of PmrA, leading to its activation as a DNA binding transcriptional regulator [68,69]. In the absence of activating signals, PmrA acts as a phosphatase that dephosphorylates (inactivates) PmrB [70]. Although genes with various functions have been identified to be part of the PmrA regulon, the predominant function of the PmrAB system appears to be controlling the expression of genes involved in the chemical modification of lipopolysaccharide (LPS) [67]. LPS modifications can have a profound impact on the nature of the Gram-negative bacterial cell envelope that, in turn, can have a range of biological effects including enabling pathogens to evade the host immune system and heightening resistance to antimicrobial compounds [71]. PmrAB is well integrated into the global regulatory network-including regulatory and functional connections with TCS such as PhoPQ and QseBC-and it is broadly distributed throughout the Enterobacterales order, including all nine of the reference strains used in this study (Fig 1C) [72,73]. However, despite its broad distribution, regulatory integration, and important function, PmrAB stood out in the analyses above as the TCS with the most variable sequence (Table 1), and as having a different divergence pattern across Enterobacteriaceae than other TCS (Fig 3C-D, S1). When compiling the TCS sequences, we noted that the genes neighbouring *pmrAB* varied in different lineages. However, because the genetic repertoires of our reference strains vary substantially, it was not immediately clear if the genomic location of *pmrAB* differed, or if there had been genetic flux at a conserved *pmrAB* genomic location. To clarify this issue, we mapped the genomic locations of the *pmrAB* locus in each of our reference strains onto the analogous location on the *E. coli* genome by identifying conserved genes that flank this locus on either side. This analysis revealed that *pmrAB* is encoded at three different loci in our reference strains, with each locus represented by three of the reference strains (Fig 5A). Relative to strains with a different *pmrAB* genomic location, all three *pmrAB* loci represent small genetic islets inserted between highly conserved genes. Interestingly, the genetic content of each islet differs: locus 1 is the largest islet and includes *pmrA/pmrB* in an operon with *pmrC*, which encodes an enzyme that chemically modifies lipid A, as well as three genes involved in acid resistance (the arginine decarboxylase system), locus 2 is comprised of only *pmrCAB,* and locus 3 encodes just the core *pmrAB* genes [74]. It is not clear which genomic location represents the ancestral locus, however *pmrAB* can be found at locus 3 outside of the Enterobacteriaceae family in certain members of the Enterobacterales Order, such as *Y. pestis* (Fig. 5A). Interestingly, analysis of the *pmrAB* genomic location in other Enterobacterales families identified a fourth *pmrAB* genomic location for members of the Pectobacteriaceae and Erwiniaceae families, which is also present as a compact three-gene *pmrCAB* islet inserted between conserved genes (Fig. S2).

**Fig 5.**
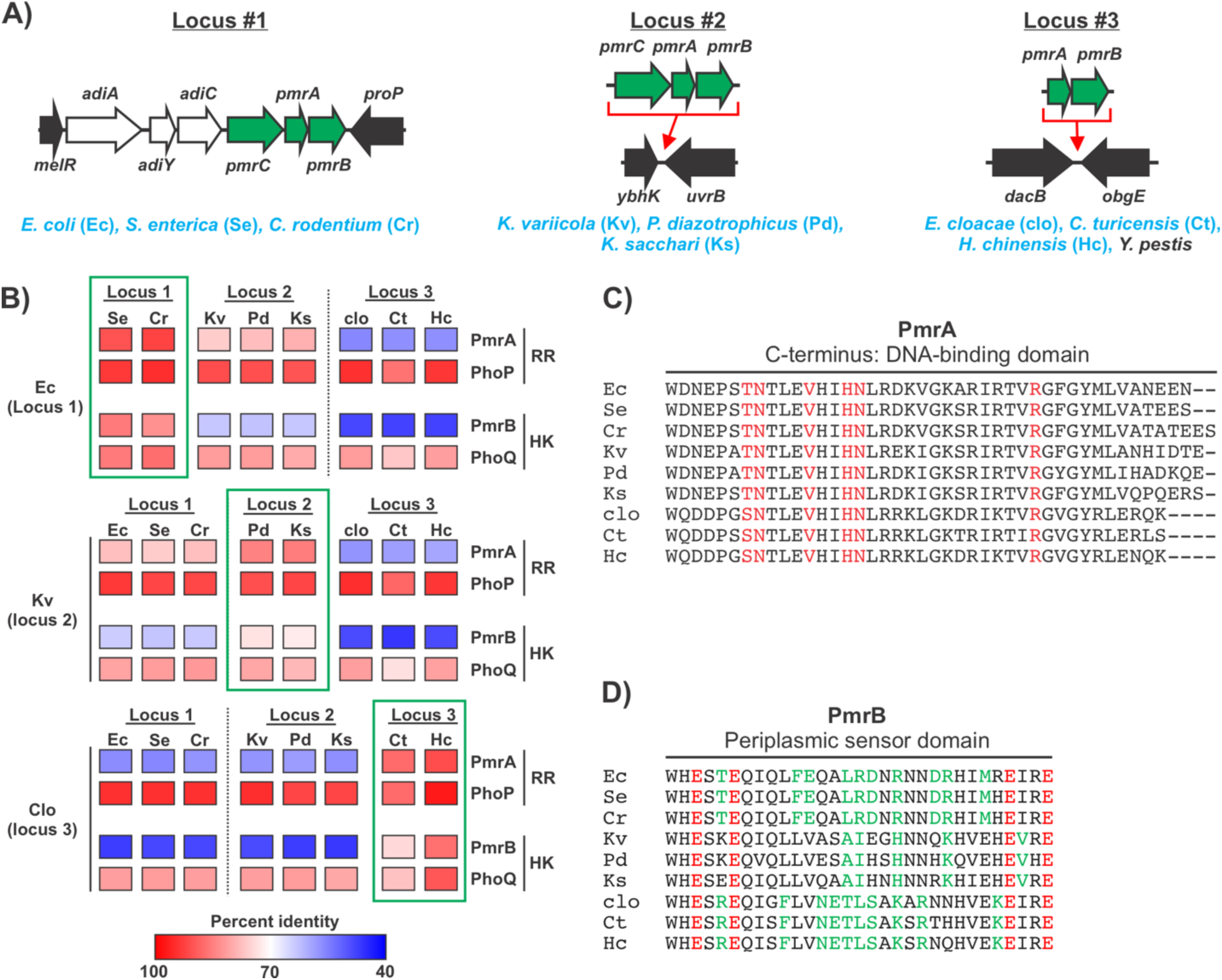
Evidence that different sequence variants of the PmrAB TCS were independently acquired by different Enterobacteriaceae lineages. **(A)** The *pmrA*/*pmrB* locus is found in three distinct genome locations amongst the strains analyzed in this study, referred to as locus 1-3. Diagram depicts these genome locations, showing their sites of insertion relative to the *E. coli* genome. Green arrows represent the *pmrC*/*pmrB*/*pmrA* genes (*pmrC* is absent from locus 3 strains), black arrows represent broadly conserved genes that flank the *pmrAB* locus that were used to map its location, and white arrows represent additional genes that are part of *pmrAB* locus 1. PmrAB is found in at locus 3 in certain species outside the Enterobacteriaceae family, including *Y. pestis*. **(B)** The unusually high level of sequence variation observed for PmrA and PmrB is only observed between species where *pmrAB* is found at different genomic locations. Diagram shows pairwise amino acid sequence comparisons (represented as heat maps) for PmrAB, as well as for a control TCS, PhoPQ. The extent of sequence variation between the RRs PhoP and PmrA and between the HKs PhoQ and PmrB is similar for comparisons where both strains carry *pmrAB* at the same genome location (green boxes). However, PmrA and PmrB show markedly more sequence variation than PhoP and PhoQ amongst strains where the *pmrAB* genome location differs (comparisons not in green boxes). **(C)** The DNA-binding amino acid residues of PmrA are highly conserved despite substantial PmrA sequence variation. Multiple amino acid sequence alignment of the C-terminal segment of PmrA that has been observed to directly contact target promoter DNA for the 9 reference strains used in this study [75]. Amino acid residues previously observed to make direct contact with the DNA of a PmrA target promoter are shown in red. **(D)** Multiple AA sequence alignment of the periplasmic sensing domain of PmrB across the reference strain set. Red resides represent the conserved “ExxE” motif that is directly involved in iron binding/sensing [68,76,77]. Green residues represent amino acid residues that are conserved amongst the PmrB encoded at the same locus, but different in all PmrB proteins found at different genomic loci. The large number of green residues suggests that the sensing properties of different PmrAB TCS might vary in a manner that correlates to genome location. Shorthand used for strains in all panels (e.g. “Ec”) is described in panel (A). Additional data relevant for this figure can be found in Fig S2.

We reasoned that if the different *pmrAB* genome locations were a result of horizontal acquisition of different alleles, that there would be a connection between the sequences of PmrA/PmrB and their genomic locations. However, such a connection would not be expected if the different locations were the result of genomic rearrangements, and the various *pmrAB* orthologs were all vertically inherited from a common ancestor. To examine this, we compared the sequence divergence of PmrAB with that of PhoPQ. PhoPQ was selected as a comparison here because it is a broadly conserved TCS that is encoded at a consistent genomic location, and which has a sequence divergence pattern that is consistent with its expected pattern based on the CG set. We found that, for both the HKs and the RRs, PmrAB and PhoPQ had similar divergence rates when comparing reference strains that have the same *pmrAB* genomic location (Fig 5B, comparisons between PhoP/PmrA and PhoQ/PmrB highlighted with green boxes). By contrast, for both the HK and the RR, PmrAB consistently exhibited markedly higher divergence rates than PhoPQ when comparing species where *pmrAB* is encoded at different genomic locations (Fig 5B, comparisons between PhoP/PmrA and PhoQ/PmrB that are not in green boxes). In accordance with this, we found that phylogenetic trees based on the sequences of PhoP or PhoQ were very similar to the tree generated based on the complete core proteome of the reference strains, consistent with PhoPQ vertical inheritance and genetic drift (Fig 1A, S2). By contrast, the phylogenetic trees generated based on PmrA and PmrB sequences clustered the reference strains into three clades based on their *pmrAB* genomic location, and the branching and branch lengths differed from the tree generated using the core proteome (Fig 1A, S2). Collectively, these data strongly suggest that the different sequence variants of the PmrAB TCS were independently acquired by different branches of the Enterobacteriaceae family.

To explore the potential functional consequences of the sequence differences amongst the various *pmrAB* loci, we compared their AA sequences focusing on their HK signal sensing domain and the RR DNA binding domain, regions directly responsible for detecting input stimuli and eliciting a biological response. To analyze PmrA’s DNA-binding motif, we took advantage of a previous study that captured the DNA-bound structure of PmrA from *K. pneumonia* [75]. We found that, despite the high levels of PmrA sequence divergence across the reference strains, all of the amino acids that directly interact with the promoter in this structure are completely conserved across all nine species (Fig 5C). This is congruent with previous observations that the promoter sequences of PmrA-regulated genes are widely conserved across different taxa [67]. Similarly, analysis of the PmrB signal sensing domain revealed that the ExxE motif required for sensing Fe^3+^ is conserved in all reference strains, in agreement with experimental data that has shown that Fe^3+^ sensing is widely conserved amongst the PmrAB systems of a wide range of species (Fig 5D) [68,76,77]. However, the amino acid sequences of the periplasmic sensing domain vary considerably amongst the representative strains, particularly between those that encode *pmrAB* at different genomic locations. We identified numerous instances of AA that are conserved amongst all reference strains with a given *pmrAB* locus, but that are not found in any reference strains whose *pmrAB* genomic location differs (Fig 5D, green residues). Importantly, we find that there is an abundance of both positively and negatively charged AA residues in this region, and that these residues vary greatly between the three different *pmrAB* loci (Fig 5D). For example, over the 31 AA periplasmic domain, there are ten AA differences that involve a charged residue between *E. coli* (locus 1) and *P. diazotrophicus* (locus 2). This remarkable variation in charged residues is significant given the essential role that ionizable AA play in sensing both Fe^3+^ and acidic pH for PmrB [68,69]. This suggests that PmrB’s sensing properties with respect to these cues (or perhaps others) is likely to differ across the Enterobacteriaceae family, particularly amongst those encoded at different genetic loci. Importantly, PmrAB plays a clinically significant role in cationic antimicrobial peptide antibiotic resistance for organisms that encode *pmrAB* at each of the three genomic locations [78–84]. Most of what is known about this TCS, however, is based on a few model species, and predominantly from studies in *S. enterica*. The data presented here suggest that caution should be taken when applying what is known about PmrAB from one species to other Enterobacteriaceae.

## DISCUSSION

In this study, we analyzed patterns of sequence change in order to investigate the evolution of TCS across the Enterobacteriaceae lineage. The factors that influence the rate of sequence change for a TCS include both positive evolutionary pressures (where mutations that alter the function of a TCS in a manner that confers a competitive advantage are more likely to become fixed in the population) and negative evolutionary pressures (where mutations that reduce the fitness of the bacterium are selected against). With respect to positive selection, higher rates of sequence change would be expected for TCS with specialized functions that vary from species to species, or for TCS that respond to signal(s) whose nature or abundance varies in a niche-specific manner. By contrast, the sequences of TCS with a stable “housekeeping” function that is conserved throughout the lineage would be expected to be less variable. With respect to negative selection, the extent to which a given TCS influences the overall fitness of the organism is likely an important factor. Even if the actions of a particular TCS are beneficial, many mutations that arise will have a very minor impact on its function, and thus their effects could be insufficient to impart a sufficient fitness disadvantage to prevent those mutations from becoming fixed. If a TCS’s sequences are highly conserved over large evolutionary distances, this therefore suggests that even minor functional changes to that TCS impart significant fitness costs. In this context, another factor that could influence how much a TCS protein diverges is the nature of the protein itself (i.e., the proportion of mutations will that influence its function). It is likely that this is a factor underlying the higher divergence rates of HKs compared to RRs, and in the weaker correlations we observe for the (architecturally diverse) sensor domain compared to other domains. However, several factors suggest this is not the predominant driving force underlying the different divergence rates of different TCS, including the strong correlations we observe between the divergence rates of the RRs and their cognate HKs, and the fact that HKs or RRs that have similar sizes, domain architectures and structures, often exhibit markedly different divergence rates. A range of other factors that relate to interactions or communication of the HK/RR pair with other regulatory factors could also influence TCS divergence rates; this would include the complexity of the TCS regulatory cascade (e.g. connector proteins, additional phosphotransfer steps, etc), the manner in which the TCS is integrated into global regulatory networks, and the need to avoid crosstalk with other TCS. Indeed, recent work has elegantly shown that the potential for crosstalk between TCS appears to have influenced the evolution of the related EnvZ/OmpR, CpxRA and RstAB TCS [38]. However, while this is undoubtedly a *bona fide* evolutionary pressure for these TCS and others, our data suggest that this is not a major driving force in the overall rate of TCS sequence change, since we found that closely related TCS that presumably face a similar pressure to avoid crosstalk can have very different divergence rates. For example, EnvZ/OmpR was the least divergent of the TCS analyzed, whereas RstAB was amongst the most divergent.

The patterns of sequence change observed in this study for different TCS or their constituent proteins or functional domains offer a window into their biology. For example, the sequence of the most conserved TCS protein, the RR OmpR, was 100% identical across all nine reference strains other than *E. cloacae* and *C. turicensis*, which have a single Ser to Ala change near the C-terminus. This extreme level of sequence conservation indicates that EnvZ/OmpR, which regulates cellular responses to osmolarity and acidic pH, plays a vital role throughout this lineage [85]. It further suggests that OmpR has a very finely tuned regulatory mechanism that is perturbed by even subtle changes to its sequence. OmpR’s regulatory mechanism is noteworthy in several ways including, (i) its recognition of target promoters exhibits limited sequence specificity and DNA structure appears to play an important role, (ii) it has been proposed that OmpR adopts different structures when bound to different target promoters in a manner that leads to differential regulation of different target genes under different conditions, (iii) its activation by EnvZ is complex has been proposed to also involve non-canonical, phosphorylation-independent activation [85–87]. Based on the data presented here, these features (and perhaps others) have imposed very stringent sequence requirements on OmpR, such that deviations from the ancestral sequence are strongly selected against and purified from the population. Interestingly, the sequence of the cytoplasmic portion of EnvZ, which is involved in both signal sensing and signal transduction, was observed to be much more variable than the rest of this TCS [66,85]. This suggests that this region might be more flexible to sequence changes without incurring perturbations to function, or that evolutionary adaptations to this TCS have been concentrated on the functions carried out by this domain.

On the opposite end of the spectrum, we find that PmrA and PmrB were the most divergent RR and HK of the proteins analyzed, and we provide evidence that this was influenced by the horizontal acquisition of different *pmrAB* alleles by different branches of Enterobacteriaceae. PmrAB has a well-established role in regulating LPS modifications, and the broad distribution of this TCS suggests that this is a function that is important throughout this family [67]. However, it is likely that the nature of the environmental conditions that would trigger the need for LPS modifications, as well as the specific response required to react to these conditions, would vary for organisms with different lifestyles and that occupy different niches. This notion is supported by studies that have identified important differences in how PmrAB is integrated into the regulatory networks of different species in this family [76,88]. Our data indicate that PmrB’s 31 AA periplasmic domain, which is well established to be essential for signal sensing in a manner that hinges on ionic interactions, is highly variable amongst different members of this family and contains a disproportionate number of charged amino acids that differ from species to species [68,69]. There is limited data available concerning how PmrB signal sensing differs in different species, but Zn^2+^ has been shown to serve as an activating signal for *E. coli* PmrB but not for *S. enterica* PmrB, indicating that its signal sensing properties are indeed evolutionarily variable [68,89]. The highly divergent PmrB sensing domains of different Enterobacteriaceae suggest that different lineages are likely to have different signal sensing properties, such as responding to different cues or responding to a given cue over a different concentration range. The sequence diversity of PmrAB in Enterobacteriaceae has direct biomedical relevance since point mutations to PmrA or PmrB are commonly observed to be the source of clinical resistance to polymyxin-family antibiotics within this family [90].

In addition to TCS-specific trends that apply across the various lineages, the data above also highlight organism-level evolutionary trends, and we identify individual TCS proteins that are more divergent than expected in a given species. In the case of *S. enterica* RstA that we highlighted above, we hypothesized that its high divergence rate might be reflective of expanded or altered functionality in this species in light of its regulation by PhoPQ (which is known to play an expanded role in *Salmonella*) and evidence that its function differs in *Salmonella* compared to other species [62]. However, the biological context underlying why specific TCS proteins have diverged more than expected in a given species could be variable. For example, in *K. variicola*, both constituents of the QseF/QseE TCS were found to be more divergent than expected. This TCS is known to be involved in regulating metabolic processes that feed into cell envelope biosynthesis, however we currently lack a complete understanding of QseEF and its cellular function [91]. While it is possible that this TCS has taken on an expanded or altered biological role in *K. variicola* that has required functional adaptations, it is also possible that this TCS is less important for fitness in this lineage and thus mutations that impact its function are less likely to be purified by natural selection. As with other findings presented here, experimental investigation will be required in order to provide functional and mechanistic context to the evolutionary trends observed.

In summary, this study has identified patterns of TCS sequence divergence across the Enterobacteriaceae family as well as individual TCS that deviate from these trends. These findings have important implications for the broader subject of how TCS evolve, and provide insight into the biology of individual TCS and their evolutionary trajectories within the Enterobacteriaceae family.

## METHODS

### Reference strains and their phylogenetic relationships

The nine reference strains selected for this study were based on previous phylogenetic analyses of the Enterobacteriaceae lineage and were selected to represent a cross section of the lineage; one strain was selected to serve as a representative of nine genera spread across this family. The selected strains were: *Salmonella enterica*, serovar Typhimurium (strain LT2), *Escherichia coli* (strain K12 MG1655), *Citrobacter rodentium* (strain ICC168), *Klebsiella variicola* (strain 342), *Cronobacter turicensis* (strain z3032), *Enterobacter cloacae* (strain ATCC 13047), *Phytobacter diazotrophicus* (strain TA9730), *Kosakonia sacchari* (strain BO-1), and *Huaxiibacter chinensis* (strain ZB04). A phylogenetic tree to analyze the evolutionary relationships between these strains was generated from whole genome sequences of the nine reference strains obtained from NCBI which were then re-annotated to extract concatenated protein sequences of single-copy core genes. Alignments of the amino acid sequences of the orthologs of all identified core genes were generated and used to infer the phylogenetic tree following the method outlined in Zheng et al [92].

### Identification and selection of TCS and CG proteins and compiling and aligning of their amino acid sequences

The 50 genes for the CG data set were selected arbitrarily on the basis that they, (i) were conserved in all reference strains, (ii) have a well-defined and conserved function, (iii) fall into diverse functional categories. AA sequences for CGs were compiled using a combination of the www.microbesonline.com sequence database and tBLASTn searches of the NCBI nr DNA sequence database using the *S*. Typhimurium protein sequences as the query [93]. The complete set of TCS encoded by both the *E. coli* and *S. enterica* reference strains was identified by generating (redundant) lists of TCS by various approaches including (i) searches of domain and protein annotations from www.microbesonline.com using these two strains (ii) analysis of the prokaryotic 2-component systems database (www.p2cs.org) for these two organisms (iii) analysis of previously published, comprehensive TCS lists for these organisms [94]. The various approaches were cross-referenced to confirm that no TCS were missed, and any inconsistencies were individually investigated. Orthologous TCS were identified in the other reference strains via tBLASTn searches using the *S. enterica* protein sequence as the query. To identify *bona fide* orthologs of RRs and HKs we set as thresholds a minimum query coverage of 90% and minimal AA percent identities of ≥60% (RRs) and ≥40% (HKs). These thresholds were set empirically on the basis that orthologous proteins (same genome location, characterized to have a similar function) were reliably captured using these cutoffs, and that TCS proteins identified by BLAST that fell below these thresholds were consistently found to be different TCS from the query sequence found at disparate genome locations. TCS that were found in fewer than 7 of the 9 reference strains were omitted from further consideration since many of the analyses conducted would be skewed by (or not interpretable for) a TCS with a narrow representation amongst our reference strains.

Once all sequences were compiled, we performed MSAs for all CGs, RRs, and HKs using Clustal Omega (default parameters) and extracted percent identity matrices for each pairwise comparison of reference strains [95]. During this process, we noted that the curated start codons often differed for certain proteins in one or more of the reference strain(s). To account for this, we manually refined our AA sequences by analyzing the DNA sequences of proteins with atypical start positions, seeking ATG, GTG, or TTG start codons that resulted in a similar sequence length to the orthologous proteins. We then repeated the MSAs for any proteins where alternative start codon selection was required.

### Generating and analyzing AA divergence rate matrices

AA sequence divergence rates (AA differences/100 AA) were generated based on the percent sequence identity matrices generated as described above. For each protein (CG, RR or HK) average AA sequence divergences rates were calculated for all pairwise combinations of reference strains. A matrix of the average of all CG values for each pairwise combinations of reference strains (“cumulative CG matrix”) was generated and used as a standard for the relative divergence rates expected of the various combinations of reference strains. To identify pairs of organisms whose RR or HK divergence rates (population level) differed from the expected values, the cumulative CG matrix was multiplied by a correction factor (HK or RR global average divergence rate/CG global average divergence rate) to obtain an expected matrix to which the observed matrix was compared. To identify individual TCS proteins whose AA sequence diverged more than expected in an individual species, the cumulative matrix was multiplied by (the average divergence rate for that protein/the CG global average) to generate an expected matrix. For any TCS protein that was absent from one or more reference strains, CG matrices and global averages used were calculated based on the same set of strains (i.e. lacking the same reference strain(s) as that TCS protein). Expected matrices were compared to observed matrices to obtain an observed/expected matrix. To determine which TCS proteins was significantly more divergent than expected, unpaired two-tailed t-tests were used comparing all observed/expected values from combinations that involve a given species to the complete set of values from combinations that do not involve that species.

### Analyzing correlations of sequence divergence rates of TCS across different lineages, of HKs and their cognate RRs, and TCS across functional domains

To analyze the extent to which TCS sequence diversity in one branch of Enterobacteriaceae correlate with the level of diversity in other branches, we selected pairs of reference stains (comparison pair) predicted to have diverged from a common ancestor after the branching point for the remaining seven reference strains, such that any sequence changes between the comparison pair are independent from any sequence changes in the remaining reference strains: *S. enterica*/*E. coli* were selected as one comparison pair and *E. cloacae*/*H. chinensis* as a second pair. Scatterplots were generated where for each TCS protein its AA sequence divergence rate for the comparison pair was plotted against its average AA sequence divergence rate for all pairwise comparisons of reference strains that do not include either strain from the comparison pair. Correlations were analyzed using the linear regression analysis tools of the OriginLab (2023b) graphing and data analysis software [96]. Outliers for this analysis were identified using a residuals plot, where proteins with a residual value greater than 3 were considered statistically significant outliers.

Analysis of the relative divergence rates of HK/RR pairs was conducted using the average values calculated across all pairwise combinations of reference strains. The values for each HK were plotted against the value for its cognate RR using a scatterplot and the linear regression analysis tools of the OriginLab (2023b) graphing and data analysis software [96]. To perform a similar analysis based on the functional domains that comprise HKs and RRs, we first parsed the amino acid sequence of each RR and HK into two functional domains. For RR, the boundaries of the receiver domain were first identified using the InterPro Protein Family Classification tool (European Bioinformatics Institute) using the AA sequence of the *S. enterica* protein as a query [97]. To ensure consistent boundaries were used, the REC domain for other reference strains were defined based on the *S. enterica* boundaries and the results of the MSA for that RR. DBD were then defined as the complete sequence of the RR that is downstream of the C-terminal boundary of the REC domain. A similar approach was used for HKs, where the AA sequence of *S. enterica* HKs were analyzed using InterPro to identify the boundaries of the kinase domain, spanning the dimerization/histidine phosphotransfer (DHp) domain combined with the ATP-binding catalytic (CA) domains. The sensor domain was then defined as the complete sequence that is upstream of the N-terminal boundary of the kinase domain, while any sequences downstream of the C-terminal boundary of the kinase domain were not included in this analysis. Domain sequences and the relationship between their divergence rates were then analyzed as described about for HK/RR pairs.

### PmrAB evolutionary analysis

The genome locations of the *pmrA* and *pmrB* (found to be adjacent in all genomes analyzed) were identified by independently analyzing the neighbouring genes in each of the reference strains. Analysis utilized the NCBI genome browser tool coupled with iterative tBLASTn searches, as well as DNA and protein-level sequence comparisons and analyses. The same approach was used to analyze *pmrAB* genome locations outside of the Enterobacteriaceae family by investigating individual strains from various families within the Enterobacterales order. Phylogenetic trees based on the sequences of PmrA, PmrB, PhoP and PhoQ were generated using the MEGA (Molecular Evolutionary Genetics Analysis) V11 software using the maximum likelihood method and a WAG +G +I substitution model based on Clustal Omega-generated MSAs. A bootstrap method with 500 total replicates was used.

## SUPPLEMENTAL INFORMATION

Supplemental information includes 2 figures, 2 tables and 6 Supplemental Datasets.

## AUTHOR CONTRIBUTIONS

The study was conceived by C.C.F. L.A.F.B, P-K.T.V. and C.C.F. contributed to experimental design and collected and analyzed the data. L.A.F.B. and C.C.F. wrote the paper with input from P-K.T.V.

## ACKNOWLEDGEMENTS

This work was supported by NSREC discovery grant (Grant number: RGPIN-2020-03964 to C.C.F.) and a start-up grant provided by the University of Alberta Faculty of Science (to C.C.F). P-K.T.V. was supported in part by an NSERC USRA award. L.A.F.B. was supported in part by an NSERC PGS-M scholarship. We would like to thank Nanzhen Qiao and the Michael Gänzle laboratory for their assistance in analyzing the phylogenetic relationships of the reference strains employed in this study.

---

**Fig. S1.**
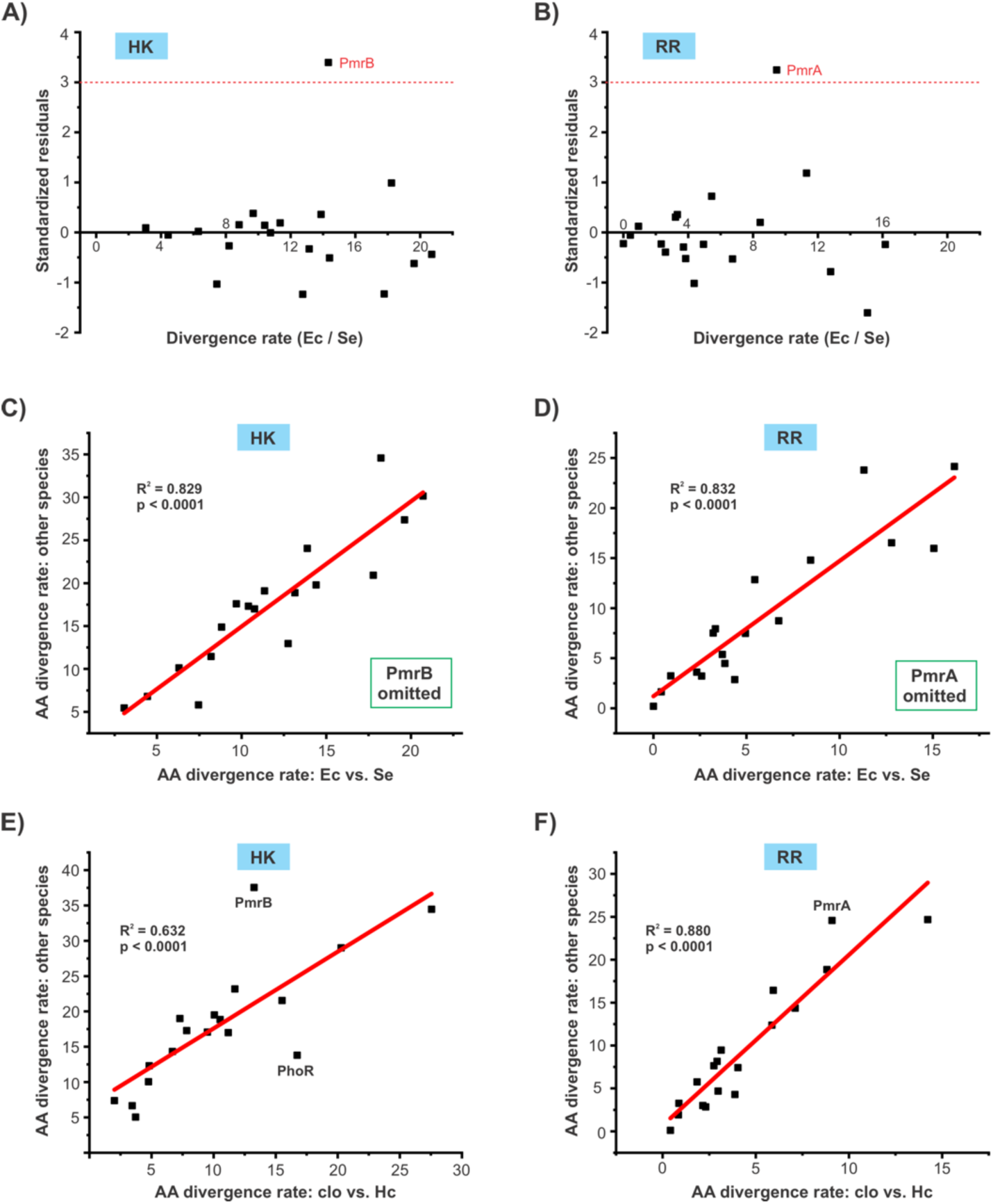
TCS sequence divergence rates between closely related genera are predictive of the divergence rates between independent Enterobacteriaceae lineages. **(A-B)** standardized residuals plots for the scatterplots presented in Fig 3C (A) and Fig 3D (B). TCS proteins with a standardized residual value >3 (red dotted line) were considered statistically significant outliers; both proteins from the PmrAB TCS were identified as outliers. **(C-D)** Scatterplots showing divergence rates (AA differences/100 AA) of each RR (A) or HK (B) for the *E. coli*/*S. enterica* pair plotted against the average rates of all pairwise combinations of the other seven Enterobacteriaceae reference strains. Plots are identical to the ones presented in Fig. 3C-D, except that the PmrAB TCS proteins were omitted because they were identified as outliers. Omitting these proteins increased the correlation coefficients of these plots substantially. **(E-F)** Scatterplots showing divergence rates (AA differences/100 AA) of each RR or HK (B) for the *E. cloacae*/*H. chinensis* pair plotted against the average rates of all pairwise combinations of the other seven Enterobacteriaceae reference strains. As with similar plots using *E. coli*/*S. enterica* as the reference pair, this comparison shows a strong positive correlation.

**Fig. S2.**
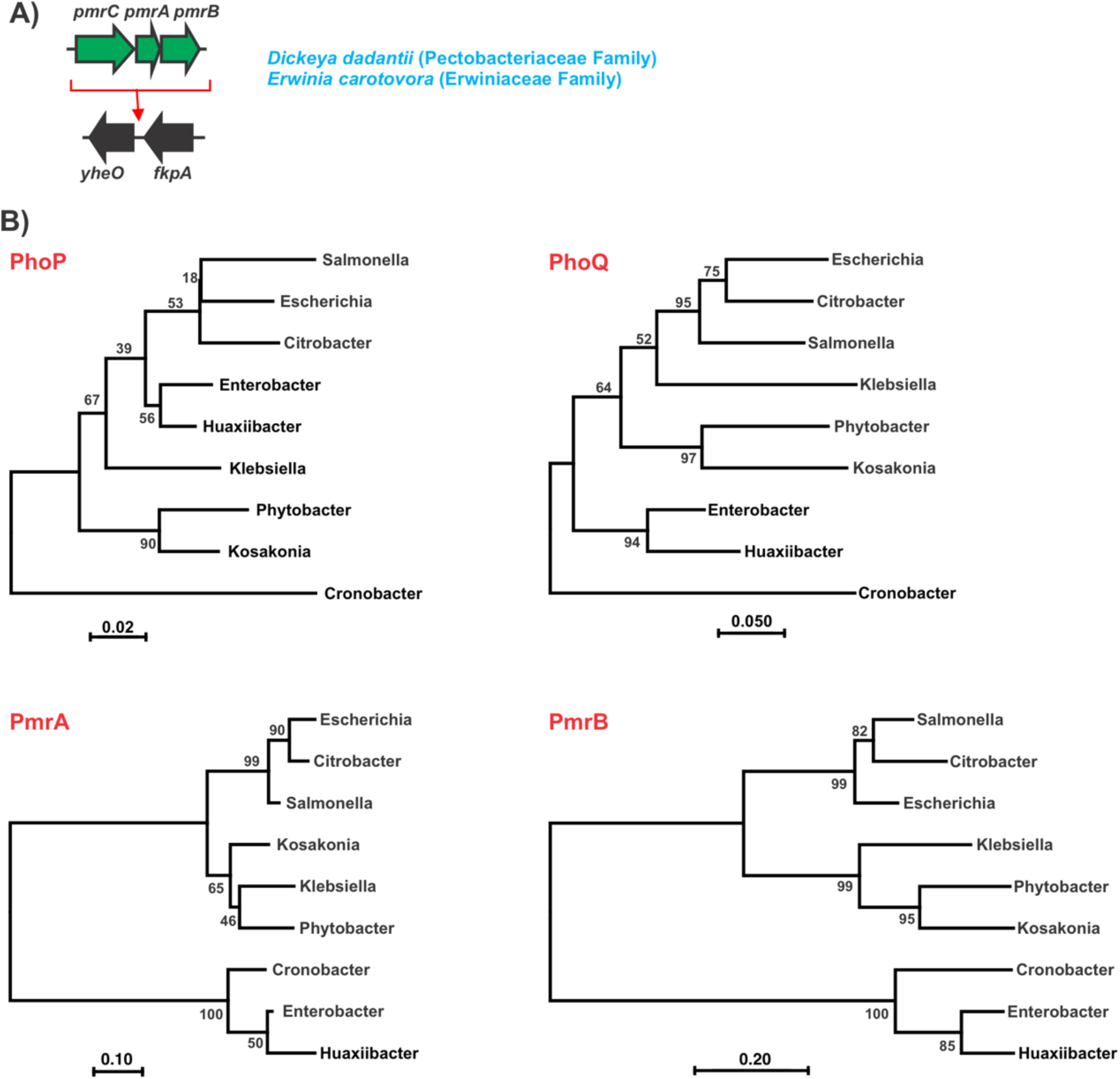
Evidence for the horizontal acquisition of PmrAB variants in different Enterobacteriaceae. **(A)** In addition to the three *pmrAB* genome locations noted for Enterobacteriaceae, it can be found in a distinct (fourth) location in certain other families of the Enterobacterales order. This genome location is shown by showing where the *pmrCAB* locus (green) is inserted relative to highly conserved genes that flank this locus on either side (black). Phylogenetic trees of the reference strains based on AA sequence alignments of the individual proteins of the PmrAB and PhoPQ TCS were generated using the MEGA (Molecular Evolutionary Genetics Analysis) V11 software using the maximum likelihood method and a WAG +G +I substitution model. A bootstrap method with 500 total replicates was used and the numbers at the nodes represent the support values. The PmrA and PmrB trees exhibit markedly different branching and branch lengths from the PhoP and PhoQ trees, as well as the tree generated based on a core proteome analysis (Fig 1A). For example, Cronobacter (*C. turicensis*) exhibits deep branching from all other strains in the trees generated for PhoP, PhoQ and the core proteome. However, Cronobacter forms a distinct clade with the other two species that carry *pmrAB* at genome locus 3 in both the PmrA and PmrB trees. Similarly, Klebsiella forms a clade with the other *pmrAB* locus 2 strains in the PmrA and PmrB trees, but not in any of the other trees.

**Table S1:** List of proteins in conserved gene (CG) data set.

| Cat* | Protein | Gene Identifier<sup>#</sup> |  |  |  |  |  | Accession Number<sup>#</sup> |  |  | Function |
| --- | --- | --- | --- | --- | --- | --- | --- | --- | --- | --- | --- |
|  |  | Se | Ec | Cr | Kv | clo | Ct | Pd | Ks | Hc |  |
| 1 | **H-NS** | STM1751 | b1237 | ROD_17711 | KPK_2107 | ECL_01633 | Ctu_23860 | BEG82436.1 | ANR77224.1 | ANG93090 | Nucleoid associated protein involved in transcriptional silencing |
| 1 | **HupB** | STM0451 | b0440 | ROD_04951 | KPK_4280 | ECL_01198 | Ctu_10200 | BEG83654.1 | ANR81039.1 | ANG91770 | Nucleoid associated protein, DNA-binding transcriptional regulator |
| 1 | **AraC** | STM0104 | b0064 | ROD_00711 | KPK_4679 | ECL_00860 | Ctu_06960 | BEG84009.1 | ANR78728.1 | ANG91513 | Arabinose operon regulatory protein |
| 1 | **SoxS** | STM4265 | b4062 | ROD_34471 | KPK_5208 | ECL_00320 | Ctu_37460 | BEG84480.1 | ANR79742.1 | ANG91143 | Superoxide response regulon transcriptional activator |
| 1 | **PurR** | STM1430 | b1658 | ROD_14111 | KPK_2351 | ECL_02341 | Ctu_19760 | BEG81792.1 | ANR77614.1 | ANG92609 | Purine nucleotide synthesis repressor |
| 1 | **Crp** | STM3466 | b3357 | ROD_44521 | KPK_0374 | ECL_04734 | Ctu_38750 | BEG79973.1 | ANR79121.1 | ANG94509 | Catabolite activator protein, cyclic AMP receptor protein |
| 1 | **Fnr** | STM1660.<br>s | b1334 | ROD_16721 | KPK_2925 | ECL_02259 | Ctu_20340 | BEG81847.1 | ANR77567.1 | ANG92667 | Fumarate nitrate reduction regulatory protein |
| 1 | **FadR** | STM1805 | b1187 | ROD_18311 | KPK_1983 | ECL_01510 | Ctu_24630 | BEG82687.1 | ANR77065.1 | ANG93138 | Fatty acid metabolism transcriptional regulator |
| 1 | **GntR** | STM3543 | b3438 | ROD_43881 | KPK_0311 | ECL_04801 | Ctu_39480 | BEG79908.1 | ANR79177.1 | ANG94575 | Gluconate utilization operon repressor |
| 1 | **DeoR** | STM0864 | b0840 | ROD_08461 | KPK_3692 | ECL_02878 | Ctu_14420 | BEG83153.1 | ANR78049.1 | ANG92188 | Deoxyribose operon repressor |
| 1 | **TreR** | STM4455 | b4241 | ROD_33121 | KPK_5015 | ECL_00659 | Ctu_36080 | BEG84253.1 | ANR79895.1 | ANG91323 | Trehalose operon repressor |
| 1 | **IhfB** | STM0982 | b0912 | ROD_09801 | KPK_3620 | ECL_02742 | Ctu_15180 | BEG83072.1 | ANR77989.1 | ANG92284 | Nucleoid associated protein, transcriptional regulation |
| 1 | **RpoS** | STM2924 | b2741 | ROD_30681 | KPK_1020 | ECL_04088 | Ctu_33130 | BEG80567.1 | ANR80230.1 | ANG93940 | RNA polymerase sigma factor |
| 1 | **ArgR** | STM3360 | b3237 | ROD_45861 | KPK_0475 | ECL_04618 | Ctu_03550 | BEG80085.1 | ANR79017.1 | ANG94400 | Arginine repressor |
| 1 | **GalR** | STM3011 | b2837 | ROD_28811 | KPK_0867 | ECL_04158 | Ctu_33870 | BEG80489.1 | ANR80191.1 | ANG94004 | Galactose operon repressor |
| 2 | **LysP** | STM2200 | b2156 | ROD_22851 | KPK_1572 | ECL_03466 | Ctu_28260 | BEG81147.1 | ANR80780.1 | ANG93507 | Lysine-specific permease |
| 2 | **BtuB** | STM4130 | b3966 | ROD_37791 | KPK_5426 | ECL_05021 | Ctu_02010 | BEG84676.1 | ANR79571.1 | ANG94740 | Vitamin B12 transporter |
| 2 | **GlnH** | STM0830 | b0811 | ROD_08121 | KPK_3729 | ECL_02920 | Ctu_14120 | BEG83204.1 | ANR78096.1 | ANG92151 | Glutamine ABC transporter |
| 2 | **ModC** | STM0783 | b0765 | ROD_07611 | KPK_3800 | ECL_02972 | Ctu_13650 | BEG83254.1 | ANR78141.1 | ANG92104 | Molybdate ABC transporter |
| 2 | **ZitB** | STM0758 | b0752 | ROD_07401 | KPK_3814 | ECL_02985 | Ctu_13520 | BEG83268.1 | ANR78156.1 | ANG92089 | Zinc transporter |
| 3 | **OmpA** | STM1070 | b0957 | ROD_10191 | KPK_3583 | ECL_02689 | Ctu_15640 | BEG82981.1 | ANR77951.1 | ANG94902 | Multifunctional and abundant outer membrane protein |
| 3 | **ExbB** | STM3159 | b3006 | ROD_35551 | KPK_0694 | ECL_04329 | Ctu_35980 | BEG80305.1 | ANR80035.1 | ANG94175 | Energetics of outer membrane transport |
| 3 | **DsbC** | STM3043 | b2893 | ROD_49401 | KPK_0777 | ECL_04219 | Ctu_34410 | BEG80433.1 | ANR80142.1 | ANG94988 | Disulfide bond formation |
| 3 | **LolA** | STM0961 | b0891 | ROD_09551 | KPK_3637 | ECL_02759 | Ctu_15030 | BEG83090.1 | ANR78000.1 | ANG92267 | Outer membrane lipoprotein carrier protein |
| 3 | **MtgA** | STM3326 | b3208 | ROD_46121 | KPK_0504 | ECL_04589 | Ctu_03710 | BEG80112.1 | ANR79001.1 | ANG94369 | Peptidoglycan biosynthesis |
| 4 | **IleS** | STM0046 | b0026 | ROD_00231 | KPK_4735 | ECL_00834 | Ctu_06520 | BEG84049.1 | ANR78753.1 | ANG91479 | Isoleucyl-tRNA synthetase |
| 4 | **EngA** | STM2519 | b2511 | ROD_24541 | KPK_1279 | ECL_03853 | Ctu_31050 | BEG80829.1 | ANR80529.1 | ANG93728 | Ribosome assembly/function |
| 4 | **SecY** | STM3420 | b3300 | ROD_45221 | KPK_0418 | ECL_04677 | Ctu_38270 | BEG80029.1 | ANR79067.1 | ANG94456 | Protein translocation |
| 4 | **RbfA** | STM3285 | b3167 | ROD_46651 | KPK_0547 | ECL_04549 | Ctu_04110 | BEG80149.1 | ANR78960.1 | ANG94319 | Ribosome maturation |
| 4 | **InfB** | STM3286 | b3168 | ROD_46641 | KPK_0546 | ECL_04550 | Ctu_04100 | BEG80148.1 | ANR78961.1 | ANG94320 | Translation initiation |
| 5 | **RecA** | STM2829 | b2699 | ROD_31151 | KPK_1096 | ECL_04033 | Ctu_32940 | BEG80637.1 | ANR80343.1 | ANG93895 | DNA recombination and repair |
| 5 | **MutS** | STM2909 | b2733 | ROD_30741 | KPK_1029 | ECL_04080 | Ctu_33050 | BEG80573.1 | ANR80232.1 | ANG93936 | DNA mismatch repair |
| 5 | **DnaC** | STM2625 | b4361 | ROD_48601 | KPK_4803 | ECL_00764 | Ctu_05720 | BEG84124.1 | ANR78810.1 | ANG91414 | DNA replication |
| 5 | **FtsA** | STM0132 | b0094 | ROD_01001 | KPK_4643 | ECL_00891 | Ctu_07270 | BEG83974.1 | ANR78703.1 | ANG91542 | Cell division process |
| 5 | **MinD** | STM1815 | b1175 | ROD_18391 | KPK_1974 | ECL_01502 | Ctu_24780 | BEG82694.1 | ANR77054.1 | ANG93143 | Septum site determination |
| 6 | **SodA** | STM4055 | b3908 | ROD_38431 | KPK_5462 | ECL_05067 | Ctu_01610 | BEG84742.1 | ANR79529.1 | ANG94787 | Oxidative stress protection |
| 6 | **DnaK** | STM0012 | b0014 | ROD_00101 | KPK_4742 | ECL_00827 | Ctu_06390 | BEG84063.1 | ANR78759.1 | ANG91473 | Chaperone protein |
| 6 | **UspA** | STM3591 | b3495 | ROD_43041 | KPK_0247 | ECL_04913 | Ctu_40120 | BEG79822.1 | ANR79237.1 | ANG94639 | Universal stress response |
| 6 | **CspD** | STM0943 | b0880 | ROD_08911 | KPK_3649 | ECL_02769 | Ctu_14900 | BEG83100.1 | ANR78012.1 | ANG92257 | Cold shock response |
| 6 | **Lon** | STM0450 | b0439 | ROD_04941 | KPK_4281 | ECL_01197 | Ctu_10190 | BEG83655.1 | ANR78450.1 | ANG91769 | Protein degradation and turnover |
| 7 | **PurF** | STM2362 | b2312 | ROD_27231 | KPK_1442 | ECL_03661 | Ctu_29400 | BEG81028.1 | ANR80691.1 | ANG93612 | Purine biosynthesis |
| 7 | **HisC** | STM2073 | b2021 | ROD_21591 | KPK_1688 | ECL_03341 | Ctu_27180 | BEG81265.1 | ANR76791.1 | ANG93389 | Histidine biosynthesis |
| 7 | **Icd (IcdA)** | STM1238 | b1136 | ROD_12261 | KPK_3407 | ECL_02498 | Ctu_17140 | BEG81619.1 | ANR77734.1 | ANG92460 | Carbon metabolism |
| 7 | **BioB** | STM0794 | b0775 | ROD_07761 | KPK_3773 | ECL_02961 | Ctu_13720 | BEG83247.1 | ANR78135.1 | ANG92115 | Biotin biosynthesis |
| 7 | **NuoC** | STM2326 | b2286 | ROD_26881 | KPK_1473 | ECL_03630 | Ctu_29110 | BEG81060.1 | ANR81212.1 | ANG93583 | Electron transport, energetics |
| 7 | **ThiL** | STM0419 | b0417 | ROD_04661 | KPK_4315 | ECL_01175 | Ctu_09940 | BEG83681.1 | ANR78472.1 | ANG91733 | Thiamine metabolism |
| 7 | **Cfa** | STM1427 | b1661 | ROD_14081 | KPK_2348 | ECL_02344 | Ctu_19730 | BEG81789.1 | ANR77617.1 | ANG92606 | Phospholipid metabolism |
| 7 | **PykF** | STM1378 | b1676 | ROD_13741 | KPK_2188 | ECL_02362 | Ctu_18790 | BEG81780.1 | ANR77621.1 | ANG92595 | Pyruvate kinase |
| 7 | **RpiA** | STM3063 | b2914 | ROD_49161 | KPK_0750 | ECL_04240 | Ctu_34630 | BEG80411.1 | ANR80124.1 | ANG94056 | Carbon metabolism |
| 7 | **ThrC** | STM0004 | b0004 | ROD_00031 | KPK_4753 | ECL_00816 | Ctu_06320 | BEG84075.1 | ANR78766.1 | ANG91461 | Threonine biosynthesis |

\* Cat indicates the functional category of the gene: 1 = gene regulation, 2 = transport, 3 = cell envelope/surface, 4 = protein synthesis, 5 = DNA replication/repair and cell division, 6 = cellular homeostasis, 7 = metabolism

**Table S2:** List of Two component system proteins included in this study.

| TCS | Gene Identifier<sup>#</sup> |  |  |  |  |  | Accession Number<sup>#</sup> |  |  | Putative function |
| --- | --- | --- | --- | --- | --- | --- | --- | --- | --- | --- |
|  | Se | Ec | Cr | Kv | clo | Ct | Pd | Ks | Hc |  |
| **PhoQ (HK)**<br>**PhoP (RR)** | STM1230<br>STM1231 | b1129<br>b1130 | ROD_12181<br>ROD_12191 | KPK_3416<br>KPK_3415 | ECL_02505<br>ECL_02504 | Ctu_17060<br>Ctu_17070 | BEG81610.1<br>BEG81611.1 | ANR77739.1<br>ANR77738.1 | ANG92455<br>ANG94907 | Mg<sup>2+</sup> homeostasis, cell surface modifications, virulence |
| **RstB (HK)**<br>**RstA (RR)** | STM1471<br>STM1475 | b1609<br>b1608 | ROD_14551<br>ROD_14561 | KPK_2940<br>KPK_2937 | ECL_02271<br>ECL_02269 | Ctu_20230<br>Ctu_20260 | BEG81835.1<br>BEG81836.1 | ANR77575.1<br>ANR80973.1 | ANG92659<br>ANG94917 | Metabolism, cell envelope biosynthesis, virulence |
| **BaeS (HK)**<br>**BaeR (RR)** | STM2130<br>STM2131 | b2078<br>b2079 | ROD_22101<br>ROD_22111 | KPK_1635<br>KPK_1634 | ECL_03405<br>ECL_03406 | Ctu_27740<br>Ctu_27750 | BEG81207.1<br>BEG81206.1 | ANR80854.1<br>ANR80853.1 | ANG93441<br>ANG93442 | Envelope stress, efflux |
| **QseC (HK)**<br>**QseB (RR)** | STM3178<br>STM3177 | b3026<br>b3025 | ROD_35681<br>ROD_35671 | KPK_0681<br>KPK_0682 | ECL_04344<br>ECL_04343 | N/A<br>N/A | N/A<br>N/A | ANR80023.1<br>ANR80024.1 | ANG94191<br>ANG94190 | Quorum sensing (?), motility, metabolism (?) |
| **EnvZ (HK)**<br>**OmpR (RR)** | STM3501<br>STM3502 | b3404<br>b3405 | ROD_44201<br>ROD_44191 | KPK_0342<br>KPK_0341 | ECL_04768<br>ECL_04769 | Ctu_39130<br>Ctu_39140 | BEG79937.1<br>BEG79936.1 | ANR79150.1<br>ANR79151.1 | ANG94542<br>ANG94543 | Osmolarity, virulence, envelope stress |
| **CpxA (HK)**<br>**CpxR (RR)** | STM4058<br>STM4059 | b3911<br>b3912 | ROD_38391<br>ROD_38381 | KPK_5460<br>KPK_5459 | ECL_05065<br>ECL_05064 | Ctu_41140<br>Ctu_41130 | BEG84738.1<br>BEG84737.1 | ANR79532.1<br>ANR79533.1 | ANG94780<br>ANG94779 | Envelope stress |
| **PmrB (HK)**<br>**PmrA (RR)** | STM4291<br>STM4292 | b4112<br>b4113 | ROD_34821<br>ROD_34831 | KPK_3766<br>KPK_3765 | ECL_04563<br>ECL_04562 | Ctu_03970<br>Ctu_03980 | BEG83240.1<br>BEG83239.1 | ANR78130.1<br>ANR78129.1 | ANG94343<br>ANG94342 | LPS modifications |
| **CreC (HK)**<br>**CreB (RR)** | STM4589<br>STM4588 | b4399<br>b4398 | ROD_50861<br>ROD_50851 | KPK_4760<br>KPK_4761 | N/A<br>N/A | Ctu_06250<br>Ctu_06240 | BEG84081.1<br>BEG84082.1 | ANR78772.1<br>ANR78773.1 | ANG91455<br>ANG91454 | Carbon metabolism |
| **NarX (HK)**<br>**NarL (RR)** | STM1766<br>STM1767 | b1222<br>b1221 | ROD_17841<br>ROD_17851 | KPK_2091<br>KPK_2089 | ECL_01620<br>ECL_01619 | Ctu_24040<br>Ctu_24050 | BEG82449.1<br>BEG82450.1 | ANR77211.1<br>ANR77210.1 | ANG93101<br>ANG93102 | Nitrogen metabolism |
| **NarQ (HK)**<br>**NarP (RR)** | STM2480<br>STM2246 | b2469<br>b2193 | ROD_24171<br>ROD_23241 | N/A<br>N/A | ECL_03766<br>N/A | N/A<br>Ctu_30620 | BEG80892.1<br>N/A | ANR80580.1<br>N/A | ANG93688<br>N/A | Nitrogen metabolism |
| **PhoR (HK)**<br>**PhoB (RR)** | STM0398<br>STM0397 | b0400<br>b0399 | ROD_04441<br>ROD_04431 | KPK_4345<br>KPK_4346 | ECL_01155<br>ECL_01154 | Ctu_09760<br>Ctu_09750 | BEG83703.1<br>BEG83704.1 | ANR81042.1<br>ANR78498.1 | ANG91712<br>ANG91711 | Phosphate metabolism |
| **KdpD (HK)**<br>**KdpE (RR)** | STM0703<br>STM0702 | b0695<br>b0694 | ROD_07011<br>ROD_07001 | KPK_3856<br>KPK_3858 | ECL_03023<br>ECL_03024 | Ctu_13110<br>Ctu_13100 | BEG83304.1<br>BEG83305.1 | ANR78186.1<br>ANR81017.1 | ANG92052<br>ANG92051 | Potassium transport |
| **GlnL (HK)**<br>**GlnG (RR)** | STM4006<br>STM4005 | b3869<br>b3868 | ROD_38921<br>ROD_38931 | KPK_5505<br>KPK_5506 | ECL_05112<br>ECL_05113 | Ctu_41850<br>Ctu_41860 | BEG84795.1<br>BEG84796.1 | ANR79490.1<br>ANR79489.1 | ANG94835<br>ANG94836 | Nitrogen metabolism |
| **DpiB (HK)**<br>**DpiA (RR)** | STM0625<br>STM0626 | b0619<br>b0620 | ROD_06341<br>ROD_06351 | N/A<br>N/A | N/A<br>N/A | N/A<br>N/A | N/A<br>N/A | N/A<br>N/A | N/A<br>N/A | SOS response, citrate/malate metabolism |
| **DcuS (HK)**<br>**DcuR (RR)** | STM4304<br>STM4303 | b4125<br>b4124 | ROD_34881<br>ROD_34871 | KPK_2904<br>KPK_2906 | ECL_03247<br>ECL_03248 | N/A<br>N/A | BEG82605.1<br>BEG82606.1 | ANR77119.1<br>ANR80928.1 | ANG94961<br>ANG93364 | Carbon<br>metabolism |
| **ArcB (HK)**<br>**ArcA (RR)** | STM3328<br>STM4598 | b3210<br>b4401 | ROD_46101<br>ROD_50881 | KPK_0502<br>KPK_4758 | ECL_04591<br>ECL_00811 | Ctu_03690<br>Ctu_06270 | BEG80110.1<br>BEG84079.1 | ANR79003.1<br>ANR78770.1 | ANG94371<br>ANG91457 | Aerobic<br>respiration<br>control |
| **TorS (HK)**<br>**TorR (RR)** | STM3826<br>STM3824 | b0993<br>b0995 | N/A<br>N/A | N/A<br>N/A | N/A<br>N/A | N/A<br>N/A | N/A<br>N/A | N/A<br>N/A | N/A<br>N/A | Anaerobic<br>respiration |
| **BarA (HK)**<br>**UvrY (RR)** | STM2958<br>STM1947 | b2786<br>b1914 | ROD_30301<br>ROD_19691 | KPK_0994<br>KPK_1869 | ECL_04118<br>ECL_01374 | Ctu_33450<br>Ctu_26160 | BEG80532.1<br>BEG81481.1 | ANR78593.1<br>ANR76897.1 | ANG93965<br>ANG93319 | Motility, stress<br>response,<br>virulence |
| **UhpB (HK)**<br>**UhpA (RR)** | STM3789<br>STM3790 | b3668<br>b3669 | ROD_40631<br>ROD_40621 | KPK_0029<br>KPK_0028 | ECL_00037<br>ECL_00036 | N/A<br>N/A | N/A<br>N/A | N/A<br>N/A | ANG90928<br>ANG90927 | Sugar uptake |
| **QseE (HK)**<br>**QseF (RR)** | STM2564<br>STM2562 | b2556<br>b2554 | ROD_24991<br>ROD_24971 | KPK_1241<br>KPK_1243 | ECL_03893<br>ECL_03891 | Ctu_31460<br>Ctu_31440 | BEG80792.1<br>BEG80794.1 | ANR80494.1<br>ANR80496.1 | ANG93766<br>ANG93764 | Metabolism of<br>cell envelope<br>precursors<br>(quorum<br>sensing?) |
| **HydH (HK)**<br>**HydG (RR)** | STM4173<br>STM4174 | b4003<br>b4004 | ROD_37491<br>ROD_37481 | KPK_5289<br>KPK_5287 | N/A<br>N/A | N/A<br>N/A | BEG84549.1<br>BEG84548.1 | N/A<br>N/A | N/A<br>N/A | Zinc transport |
| **CopS (HK)**<br>**CopR (RR)** | STM1095<br>STM1096 | b1968<br>b1969 | ROD_50431<br>ROD_50441 | N/A<br>N/A | N/A<br>N/A | Ctu_41020<br>Ctu_41030 | N/A<br>N/A | N/A<br>N/A | N/A<br>N/A | Unknown (copper<br>homeostasis?) |
| **YehU (HK)**<br>**YehT (RR)** | STM2159<br>STM2158 | b2126<br>b2125 | ROD_22391<br>ROD_22381 | KPK_1605<br>KPK_1606 | ECL_03436<br>ECL_03435 | Ctu_27900<br>Ctu_27890 | BEG81185.1<br>BEG81186.1 | ANR80816.1<br>ANR80817.1 | ANG94970<br>ANG93480 | Carbon<br>metabolism |
| **RcsC (HK)**<br>**RcsB (RR)** | STM2271<br>STM2270 | b2218<br>b2217 | ROD_23521<br>ROD_23501 | KPK_1512<br>KPK_1513 | ECL_03522<br>ECL_03521 | Ctu_28750<br>Ctu_28740 | BEG81098.1<br>BEG81099.1 | ANR80742.1<br>ANR80743.1 | ANG93545<br>ANG93544 | Cell envelope<br>stress |
| **CheY (HK)**<br>**CheA (RR)** | STM1916<br>STM1921 | b1882<br>b1888 | ROD_19301<br>ROD_19411 | N/A<br>N/A | ECL_1404<br>ECL_1416 | Ctu_25740<br>Ctu_25850 | BEG82865.1<br>BEG82853.1 | ANR76915.1<br>ANR76926.1 | ANG93288<br>ANG93276 | Chemotaxis |

#: Acronyms for strains for gene identifiers or accession numbers are as follows: Se = *Salmonella enterica* (serovar Typhimurium, strain LT2), Ec = *Escherichia coli* K-12 (strain MG1655), Cr = *Citrobacter rodentium* (strain ICC168), Kv = *Klebsiella variicola* (strain 342), Ct = *Cronobacter turicensis* (strain z3032), clo = *Enterobacter cloacae* (strain ATCC 13047), Pd = *Phytobacter diazotrophicus* (strain TA9730), Ks = *Kosakonia sacchari* (strain BO-1), and Hc = *Huaxiibacter chinensis* (strain ZB04)

